# Liquid-like condensates that bind actin drive filament polymerization and bundling

**DOI:** 10.1101/2024.05.04.592527

**Authors:** Caleb Walker, Aravind Chandrasekaran, Daniel Mansour, Kristin Graham, Andrea Torres, Liping Wang, Eileen M. Lafer, Padmini Rangamani, Jeanne C. Stachowiak

**Author notes:** These authors contributed equally (Aravind Chandrasekaran, Daniel Mansour).

## Abstract

Liquid-like protein condensates perform diverse physiological functions. Previous work showed that VASP, a processive actin polymerase, forms condensates that polymerize and bundle actin. To minimize their curvature, filaments accumulated at the inner condensate surface, ultimately deforming the condensate into a rod-like shape, filled with a bundle of parallel filaments. Here we show that this behavior does not require proteins with specific polymerase activity. Specifically, we found that condensates composed of Lamellipodin, a protein that binds actin but is not an actin polymerase, were also capable of polymerizing and bundling actin filaments. To probe the minimum requirements for condensate-mediated actin bundling, we developed an agent-based computational model. Guided by its predictions, we hypothesized that any condensate-forming protein that binds actin could bundle filaments through multivalent crosslinking. To test this idea, we added an actin-binding motif to Eps15, a condensate-forming protein that does not normally bind actin. The resulting chimera formed condensates that drove efficient actin polymerization and bundling. Collectively, these findings broaden the family of proteins that could organize cytoskeletal filaments to include any actin-binding protein that participates in protein condensation.

## Introduction

The actin cytoskeleton forms filament networks that play a critical role in cell motility, endocytosis, and adhesion.^1–4^ The assembly of these higher-order structures is facilitated by a family of actin accessory proteins, which collectively determine filament polymerization rate, length, and arrangement into networks.^5–7^ Several cytoskeletal accessory proteins have recently been shown to assemble into condensates via liquid-liquid phase separation (LLPS), a mechanism by which biomolecules self-assemble into a liquid-like condensed phase surrounded by a dilute phase.^8–10^ Interestingly, these biomolecular condensates can nucleate the assembly of cytoskeletal filaments. For example, condensates consisting of proteins from the T-cell receptor phosphorylation cascade are capable of concentrating and polymerizing actin.^11^ Similarly, condensates formed from the *C. elegans* tubulin polymerase SPD-5 can nucleate microtubule aster formation in the presence of the microtubule-stabilizing proteins TPXL-1 and ZYG-9.^12^ Building on these findings, we recently showed that condensates consisting of the actin polymerase VASP can polymerize and bundle actin filaments.^13^ As a homo-tetramer with a high degree of intrinsic disorder, VASP has key hallmarks of proteins that form biomolecular condensates.^14–16^ As actin polymerized inside VASP condensates of micrometer diameter, actin filaments, which have a persistence length of 10-20 micrometers,^17^ accumulated at the inner surfaces of condensates to minimize filament curvature. This partitioning led to the assembly of a peripheral, ring-like bundle of actin within condensates. As actin continued to polymerize, an increasing number of filaments joined this ring, increasing its rigidity. When the rigidity of the actin ring overcame the surface tension of the VASP condensate, the filaments within the ring began to straighten, deforming the initially spherical VASP condensates into elliptical and rod-like structures filled with parallel bundles of actin filaments.^13,18^

In cells, VASP works together with multiple other cytoskeletal accessory proteins to drive filament polymerization and bundling. Each monomer of VASP consists of an N-terminal EVH1 domain, which binds to short proline-rich sequences in its binding partners. The EVH1 domain is followed by VASP’s central proline-rich region, and then by an EVH2 domain through which VASP binds and polymerizes actin. Finally, VASP contains a c-terminal tetramerization domain.^14–16^ The EVH1 domain of VASP interacts with proline-rich repeats in multiple cytoskeletal accessory proteins, many of which are native multimers with a high degree of intrinsic disorder.^19–22^ These features suggest that VASP’s binding partners could reinforce its condensation, helping to build a more stable protein network that is capable of polymerizing actin and controlling the morphology of the resulting filament network. As one example, our recent work showed that the addition of Arp2/3, which nucleates the assembly of branched actin networks, to VASP condensates results in aster-shaped structures. This observation illustrates that condensates of differing compositions can facilitate varying actin network morphologies.^23^ Another VASP binding partner is Lamellipodin, which has been shown to interact with VASP in cytoskeletal protrusions such as lamellipodia and filopodia.^20,24–27^ Lamellipodin dimerizes via an N-terminal coiled-coil domain,^28^ which is followed by a Ras-associating and Pleckstrin homology domain (RA-PH), which allows Lamellipodin to localize to the plasma membrane via lipid binding.^29^ After the RA-PH domain Lamellipodin’s c-terminus is proline-rich and almost completely disordered. It is within this c-terminal disordered region that Lamellipodin contains several proline-rich regions that include multiple EVH1 binding sequences.^20,26^ While Lamellipodin binds actin filaments,^25,30^ it lacks actin polymerase activity.^25^ This observation has led to the suggestion that Lamellipodin’s role is to recruit and cluster other cytoskeletal accessory proteins, such as VASP, during membrane remodeling events.^25^ Interestingly, Lamellipodin and VASP form dynamic clusters at the leading edge of motile cells, which undergo dynamic fission and fusion events.^24,25^ This observation, in addition to the recent finding that VASP and Lamellipodin co-partition into protein condensates,^31^ suggests that the two proteins form a flexible, liquid-like network.

Here we asked how interactions between VASP and Lamellipodin impact the ability of protein condensates to polymerize and bundle actin filaments. We began by characterizing the ability of Lamellipodin to form liquid-like condensates *in vitro* and to stabilize the assembly of VASP condensates. When actin was added to these condensates, it polymerized and formed bundles, deforming the condensates into rod-like structures, similar to our previous observations with condensates consisting of VASP alone.^13^ Surprisingly, we found that condensates consisting of Lamellipodin alone can also polymerize and bundle actin filaments, despite Lamellipodin’s reported lack of polymerase activity. How does the formation of protein condensates confer this capacity upon Lamellipodin? Multi-valent binding to actin filaments is thought to underlie the ability of specialized actin polymerases, such as formins and members of the ENA/VASP family, to stabilize filament bundles and add monomers to growing filament nuclei ^7,14,32–34^ Therefore, one possible explanation for the ability of Lamellipodin condensates to polymerize and bundle actin is that the condensate environment promotes multivalent interactions between Lamellipodin and actin. To investigate the potential contribution of protein condensates to actin bundling, we developed an agent-based model of filament rearrangement within spherical containers that mimic protein condensates. In this context, we examined the ability of actin accessory proteins, such as VASP and Lamellipodin, to bundle actin filaments. This model predicted that any actin-binding protein that forms a multi-valent complex, either stably or dynamically, is sufficient to bundle actin filaments. From this perspective, virtually any actin-binding protein that forms condensates should be able to drive actin polymerization and bundling. To test this hypothesis, we formed protein condensates of Eps15, an endocytic protein lacking known interactions with actin.^35–37^ When we added the actin-binding motif, Lifeact, to the C-terminus of this protein, the resulting chimera formed condensates that spontaneously polymerized and bundled actin filaments. Collectively, these results suggest that filament polymerization and bundling are emergent properties of liquid-like protein condensates that bind actin. Given that many actin-interacting proteins are now thought to participate in condensates,^11,13,23,38–42^ our results suggest a general principle of actin organization through multivalent interactions.

## Results

### Lamellipodin phase separates into liquid-like condensates

As a native dimer with a high degree of intrinsic disorder, Lamellipodin has key hallmarks of the ability to form biomolecular condensates.^8,9^ In particular, its largely disordered C-terminal region (residues 850-1250) contains twenty-seven negatively charged residues (aspartate and glutamate) and forty-four positively charged residues (lysine and arginine)^20,25^ suggesting a strong potential for intra and inter-molecular electrostatic interactions. Further, the same region contains ninety-nine proline residues, which tend to increase chain rigidity, contributing to intrinsic disorder and phase separation.^43,44^ To study the potential of Lamellipodin to phase separate *in vitro*, we used a minimal model of the full-length protein for ease of expression and purification, as has been previously reported.^24,25^ This minimal protein, which we will refer to as mini-Lpd, consisted of an N-terminal GFP domain, followed by a dimerizing leucine zipper motif to imitate native dimerization, and ending with the c-terminal disordered region of Lamellipodin, residues 850-1250 **(Fig. 1A,B)**. To test the ability of mini-Lpd to form phase-separated protein condensates, we mixed 5 - 15 μM mini-Lpd with 3% (w/v) PEG 8000. PEG is commonly added in the study of LLPS to mimic the crowded environment in the cell cytoplasm.^45–47^ Upon the addition of PEG, mini-Lpd formed spherical protein condensates with diameters in the micrometer range, which increased in size with increasing protein concentration **(Fig. 1C)**. These condensates fused and re-rounded upon contact within a second (**Fig. 1D),** suggesting liquid-like behavior.^10,48^ Additionally, mini-Lpd condensates recovered rapidly after photobleaching, indicating dynamic molecular exchange **(Fig. 1E,F).** To explore the contribution of electrostatic interactions to the condensation of mini-Lpd, we examined condensates in buffers of increasing ionic strength **(Fig. 1G).** The average size of condensates, as well as protein partitioning into them, decreased substantially as ionic strength increased, suggesting that they were stabilized by electrostatic interactions, which are screened at high ionic strength (**Fig. 1H,I)**.

**Figure 1.**
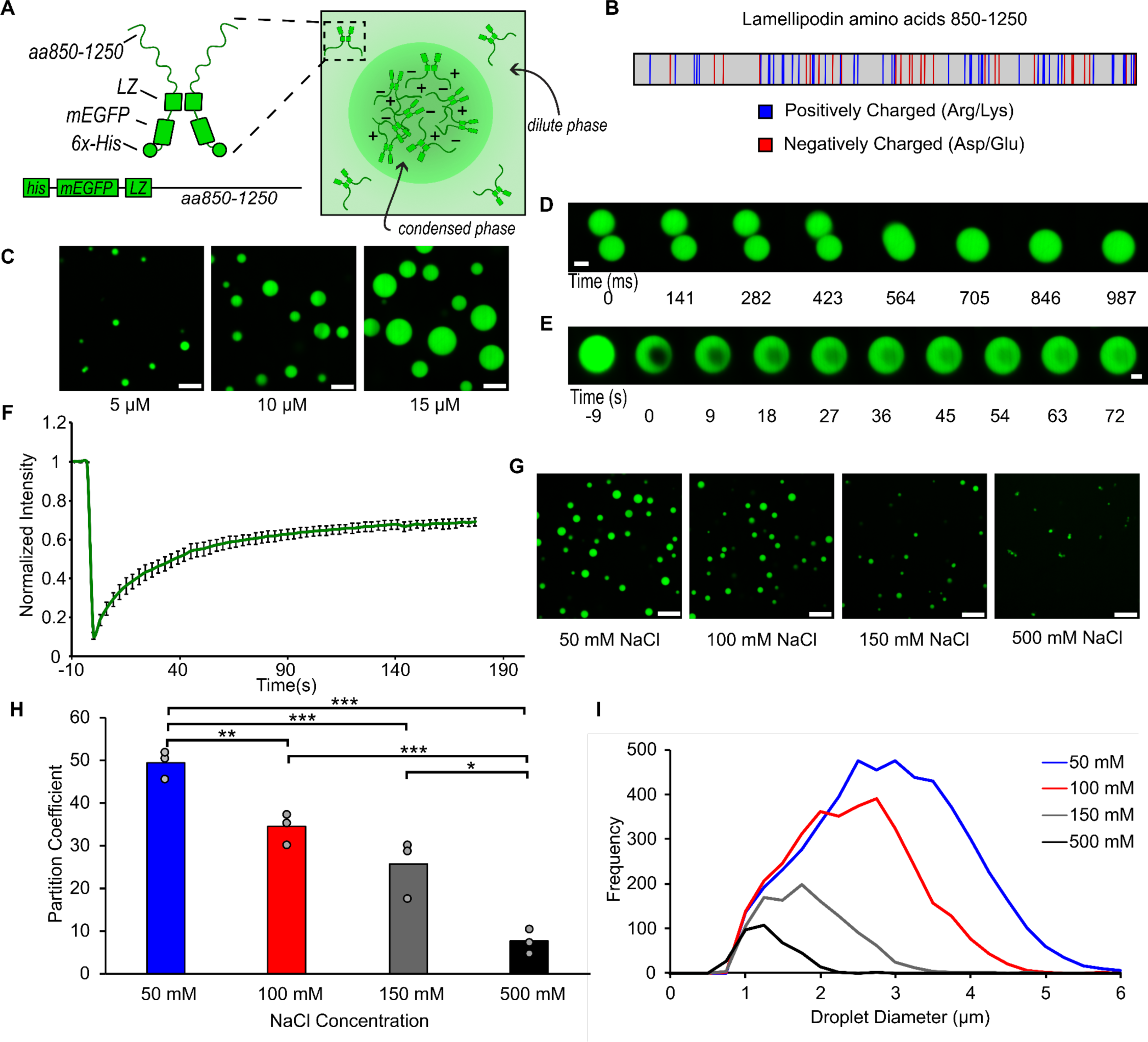
A minimal version of Lamellipodin phase separates into liquid-like condensates. **A)** Left: Schematic depicting domains of mini-Lpd (LZ: Leucine Zipper). Right: schematic depicting condensate formation **B)** Amino acid sequence of amino acids 850-1250 of Lamellipodin with positively charged (blue) and negatively charged (red) amino acids highlighted. **C)** Condensates formed by mini-Lpd at increasing protein concentrations in a buffer containing 20 mM Tris (pH 7.4), 150 mM NaCl, and 5 mM TCEP. Scale bars are 5 μm. **D)** Time course of condensate fusion event for 10 μM mini-Lpd in buffer containing 20 mM Tris (pH 7.4), 50 mM NaCl, and 5 mM TCEP. Scale bar 2 μm. **E)** Representative images of fluorescence recovery after photobleaching of a mini-Lpd condensate. Scale bar 2 μm. **F)** Plot of average fluorescence recovery after photobleaching for mini-Lpd condensates. Lines are the average recovery +/- s.d. at each timepoint for each protein across n=6 independent samples. **G)** mini-Lpd condensates formed in buffers with increasing ionic strength. mini-Lpd concentration is 10 μM in all conditions. Scale bars are 5 μm. **H)** Quantification of the partitioning of mini-Lpd into condensates formed from 10 μM mini-Lpd under the conditions shown in **G.** Partition coefficient is defined at the ratio of protein intensity inside the condensates to that in the bulk solution. Bars represent the average across three independent experiments with at least three images quantified per experiment. One asterisk denotes p<.05, two asterisks denote p<.01, and three asterisks denote p<.001 using an unpaired, two-tailed t-test on the means of the replicates N=3. **I)** Distribution of condensate diameters for the conditions shown in **G** across three separate replicates for each condition.

### Interactions between Lamellipodin and VASP mutually stabilize protein condensation

Having established that mini-Lpd can undergo LLPS, we next investigated the potential impact of mini-Lpd on the phase separation of VASP. Our previous work showed that VASP forms liquid-like condensates across a range of protein concentrations and ionic strengths when 3% (w/v) PEG 8000 is used as a crowding agent.^13^ We hypothesized that adding mini-Lpd could strengthen the VASP network by forming multivalent interactions between the multiple proline-rich motifs in each Lamellipodin protein, which are recognized by the four EVH1 domains in the VASP tetramer **(Fig. 2A).** To test this idea, we first confirmed that neither VASP nor mini-Lpd were able to form condensates in the absence of PEG **(Fig. 2B).** We then combined mini-Lpd and VASP at increasing ratios, keeping the total protein concentration constant at 30 μM. Beginning with pure mini-Lpd, we gradually increased the VASP concentration from 7.5 μM to 22.5 μM, finding a range of ratios from 3:1 mini-Lpd:VASP to 1:3 mini-Lpd:VASP for which condensate formation was observed in the absence of PEG crowding **(Fig. 2C,D).** Condensates of mini-Lpd and VASP that formed in the absence of PEG retained liquid-like behaviors, merging and re-rounding upon contact in less than a second **(Fig. 2E)**, and recovering rapidly after photobleaching **(Fig. 2F,G**). These results indicate that mini-Lpd and VASP form an interconnected network that condenses in the absence of exogenous crowding agents. To test the dependency of this network on multi-valent contacts between VASP and mini-Lpd, we evaluated mutants of VASP that would be expected to inhibit such contacts. These included a monomeric version of VASP (mVASP), which lacked the tetramerization domain, and a version of VASP lacking the EVH1 domain, ΔEVH1-VASP. Both proteins failed to form condensates upon the addition of mini-Lpd at concentrations that drove condensation of wild-type VASP. These data suggest that multivalent contacts between VASP and mini-Lpd are essential to the co-condensation of the two proteins **(Fig. 2H,I).**

**Figure 2.**
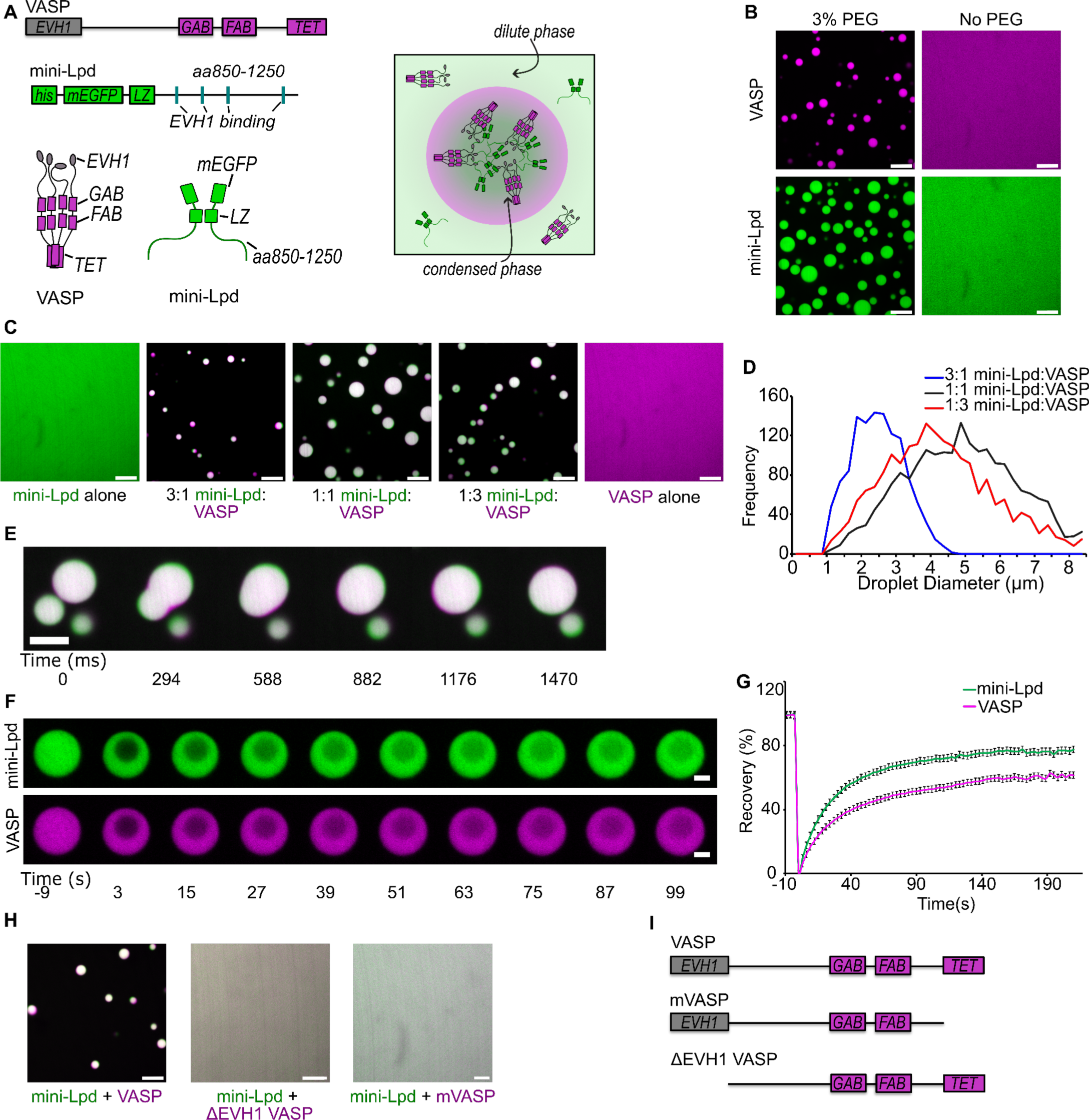
Interactions between Lamellipodin and VASP mutually stabilize protein condensation. **A)** Top: Schematic of domain organization in a VASP monomer (GAB: G-actin binding site; FAB: F-actin binding site; TET: Tetramerization domain) and a mini-Lpd monomer (LZ: Leucine Zipper dimerization motif) Bottom: Schematic of VASP tetramer, mini-Lpd dimer, and condensate formation with both proteins present. Right: Schematic depicting mini-Lpd and VASP co-partitioning into a condensate. **B)** Condensates form upon the inclusion of 3% w/v PEG in solution for both VASP (magenta) and mini-Lpd (green); however, neither protein forms condensates in buffer lacking PEG. Scale bars 5 μm. Protein concentrations were 20 μM for both the with and without PEG conditions. **C)** Panels showing representative images of mini-Lpd + VASP condensates formed at various mini-Lpd to VASP ratios. Scale bars 5 μm. **D)** Distribution of condensate diameters for each condition in **C**.**E)** Time course of a condensate fusion event between mini-Lpd (green) and VASP (magenta) condensates. Scale bar 2 μm. **F)** Representative images of fluorescence recovery after photobleaching of a mini-Lpd (green) and VASP (magenta) condensate. Scale bars 2 μm. **G)** Plot of average fluorescence recovery after photobleaching for mini-Lpd and VASP condensates formed in the absence of PEG. Lines are the average recovery +/- s.d. at each timepoint for each protein across n=9 independent samples. **H)** mini-Lpd added to the respective VASP mutant in a 1:1 ratio to test for condensate formation in the absence of PEG. Left – mini Lpd (green) and VASP (magenta), Middle - mini-Lpd (green) and VASPΔEVH1 (magenta), Right - mini-Lpd (green) and monomeric VASP (mVASP) (magenta). Scale bars 5 μm **I)** Diagrams depict domain structures of VASP mutants. All experiments were performed in a buffer containing 20 mM Tris (pH 7.4), 150 mM NaCl, and 5 mM TCEP in the absence of PEG, except where noted in panel **B** where 3% (w/v) PEG was included.

### Condensates of VASP and mini-Lpd polymerize and bundle actin

Next, we evaluated the ability of VASP/mini-Lpd condensates to polymerize and bundle actin filaments. We added increasing concentrations of monomeric actin (G-actin) to protein condensates formed from mini-Lpd and VASP in the absence of PEG. As the concentration of actin increased, the protein condensates began to deform, taking on increasingly elongated shapes **(Fig. 3A)**. We confirmed actin polymerization within the condensates by phalloidin staining, which specifically binds to filamentous actin (**Fig. 3B)**, permitting us to visualize the gradual deformation of condensations from initially spherical shapes to ellipsoids and finally to rod-like geometries **(Fig. 3C,D)**. These morphological changes are in line with our previous observations with condensates consisting of VASP alone.^13^ For condensates consisting of VASP and mini-Lpd, the aspect ratio (longest dimension divided by shortest dimension) of the condensates increased with increasing actin concentration, as did the fraction of condensates with aspect ratios above a threshold value of 1.2 (**Fig. 3E,F)**. Together these data indicate that condensates of VASP and mini-Lpd, which form without the requirement for crowding agents, drive local polymerization and bundling of actin filaments.

**Figure 3.**
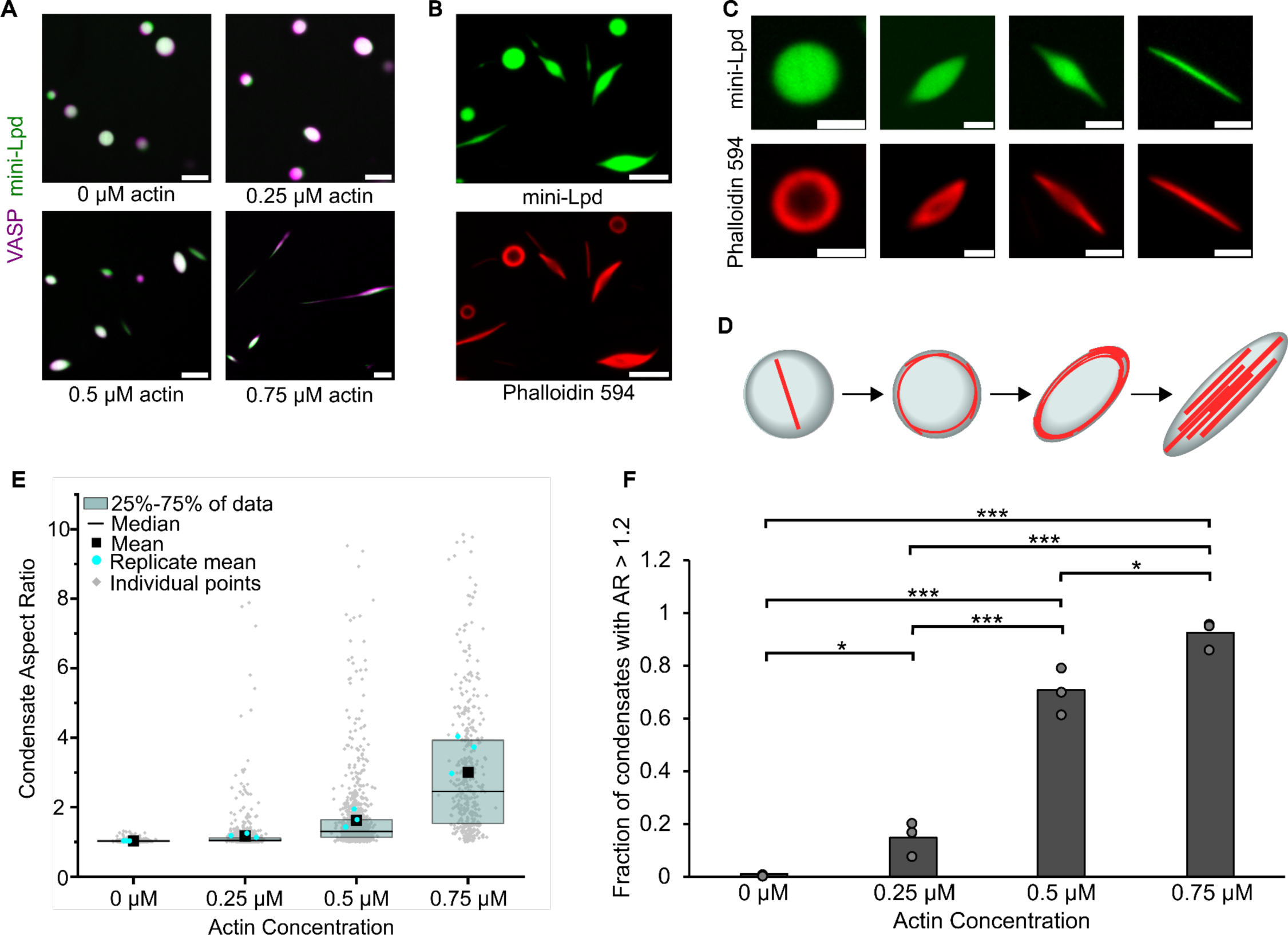
Condensates of VASP and mini-Lpd polymerize and bundle actin in the absence of crowding agents. **A)** Condensates formed from 5 μM mini-Lpd (green) and 10 μM VASP (magenta) are increasingly deformed with the addition of increasing concentrations of G-actin (unlabeled). Scale bars 5 μm **B)** Phalloidin-iFluor-594 (red) staining of mini-Lpd (green) and VASP (unlabeled) condensates with 0.5 μM monomeric G-actin displaying rings and rods of polymerized actin within the protein condensates. Scale bars 5 μm. **C)** From left to right: representative confocal images depicting the progression of condensate deformation as actin polymerizes and bundles within the protein condensates. Scale bars 2 μm **D)** Cartoon depicting the mechanism of actin polymerization within protein condensates and the role it plays in condensate deformation. **E)** Distribution of condensate aspect ratios across the conditions in **A**, with at least 400 condensates analyzed for each condition, and more than 800 condensates were analyzed for the 0, 0.25, and 0.5 μM conditions. In the 0.75 μM actin condition values for aspect ratios above 10, corresponding to 4.8% of the data, are not displayed to better visualize distributions for all conditions. The mean indicates the mean of all data points in each condition, while the replicate mean is the mean of each of the individual replicates for each condition. **F)** Quantification of the fraction of elongated protein condensates, defined as condensates with aspect ratios > 1.2, across the conditions in **A**. Data are mean across three independent experiments with at least 400 condensates analyzed per condition, and more than 800 condensates were analyzed for the 0, 0.25, and 0.5 μM conditions. Overlayed gray circles denote the means of each replicate. One asterisk denotes p<.05, three asterisks denote p<.001 using an unpaired, two-tailed t-test on the means of the replicates N=3. All experiments were performed in a buffer containing 20 mM Tris (pH 7.4), 150 mM NaCl, and 5 mM TCEP in the absence of PEG.

### Condensates of mini-Lpd polymerize and bundle actin in the absence of VASP

VASP is a well-characterized actin polymerase and bundling protein.^14,32,49^ In contrast, Lamellipodin binds actin filaments but fails to increase their rate of barbed end elongation.^25^ Therefore we expected that condensates formed from mini-Lpd alone **(Fig. 1)** would fail to polymerize and bundle actin. To test this assumption, we added monomeric actin to pre-formed condensates of mini-Lpd. Surprisingly, we found that the condensates deformed upon actin addition, suggesting that actin was polymerizing and forming bundles inside the condensates **(Fig. 4A).** When Latrunculin A, an inhibitor of actin polymerization,^50^ was added to the condensates prior to actin addition, it inhibited both actin polymerization and condensate deformation, establishing that actin polymerization led to deformation of mini-Lpd condensates **(Fig. 4B).** Increasing the concentration of actin added to mini-Lpd condensates resulted in higher aspect ratios and a higher fraction of condensates with an aspect ratio above 1.2 **(Fig. 4C,D)**. Phalloidin staining revealed that filamentous actin bundles were present inside mini-Lpd condensates as they deformed from spherical to rod-like structures **(Fig. 4E-F).** Owing to these surprising results, we decided to perform simulations aimed at determining the key physical requirements for filament bundling by protein condensates.

**Figure 4:**
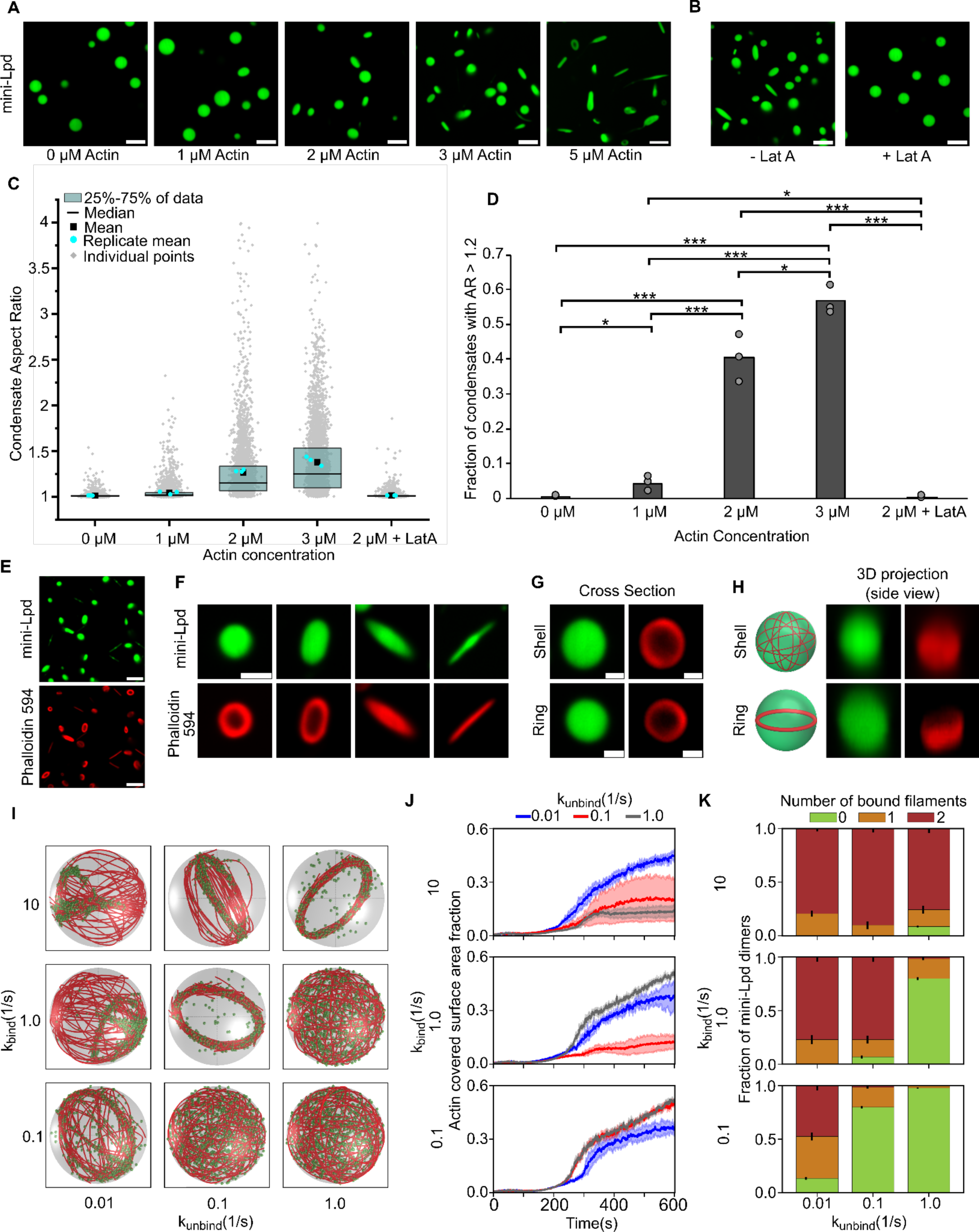
Condensates of mini-Lpd polymerize and bundle actin. **A)** The addition of actin at increasing concentrations to condensates formed from 5 μM mini-Lpd (green) results in increasingly deformed protein condensates. Scale bars 5 μm. **B)** Pretreatment of mini-Lpd condensates with 5 μM latrunculin A (LatA) prior to G-actin addition inhibits actin polymerization and results in spherical condensates. Scale bars 5 μm. **C)** Distribution of condensate aspect ratios across the conditions in **A**, with at least 1000 condensates analyzed for each condition. The mean is the mean of all data points in each condition, while the replicate mean is the mean of each of the individual replicates for each condition. **D)** Quantification of the fraction of high-aspect-ratio protein condensates, defined as condensates with aspect ratios > 1.2, across the conditions in **A**. Data are mean across three independent experiments with at least 1000 condensates analyzed per condition. Overlayed gray circles denote the means of each replicate. One asterisk denotes p<.05, two asterisks denote p<.01, and three asterisks denote p<.001 using an unpaired, two-tailed t-test on the means of the replicates, N=3. **E)** Phalloidin-iFluor-594 (red) staining of mini-Lpd condensates (green) with 5 μM monomeric G-actin (unlabeled) added displaying rings and rods of polymerized actin within the protein condensates. Scale bars 5 μm. **F)** Representative confocal cross-section images of the progression of condensate deformation as a result of actin polymerization. Scale bar 1 μm. **G)** Representative 2D confocal images of independent mini-Lpd condensates (green) containing peripheral actin, shown with phalloidin staining (red), in a shell of actin (top) and a ring of actin (bottom). Scale bars 1 μm. **H)** 3D reconstructions of the same mini-Lpd condensates shown in **G** demonstrating a shell of actin (top) and a ring of actin (bottom). **I)** Simulations show that bivalent crosslinker kinetics affect actin network organization in LLPS condensates. Representative final snapshots (t = 600 s) from simulations at various binding and unbinding rates within spherical condensates (R = 1 μm) containing 30 actin filaments (red) and 1000 bivalent crosslinkers (green spheres). Please refer to the Supplementary Methods section for a detailed description of the model. The binding rates of the bivalent crosslinkers are varied along each column, and unbinding rates are varied along each row. The polymerization rate at the plus (+) end is constant at 0.0103 μm/s, and neither end undergoes depolymerization. See also **Supplementary Movie M2**. For additional kinetic conditions, see **Figure S4** and **Supplementary Movie M3**. **J)** Dynamics of the actin-covered surface area fraction for varied bivalent crosslinker binding and unbinding kinetics. (Data used: 10 replicates) **K)** Stacked bar graphs representing the fraction of bivalent crosslinkers bound to 0, 1, or 2 actin filaments for each condition. The error bars represent the standard deviation. For **J** and **K**, 10 replicates are considered per condition, and the data was obtained from the last 30 snapshots (5%) of each replicate. For analysis of additional kinetic conditions, see **Figure S5**.

### Agent-based simulations predict that multivalent cross-linking of actin filaments drives condensate-mediated bundling

Our previous work showed that the formation of a ring-like bundle of actin filaments within condensates is a critical step along the path to condensate deformation by actin.^13,18^ Specifically, as actin filaments polymerized inside condensates, they partitioned to the inner surface of the condensate to minimize their curvature, creating a shell of actin filaments. As this three-dimensional shell formed, it gradually collapsed into a two-dimensional ring-like bundle **(Fig. 4G,H)**. This collapse concentrated pressure on the condensate, overcoming its surface tension such that filaments straightened into a parallel, rod-like geometry.^13,18^ We sought to understand how actin-binding proteins such as VASP and Lamellipodin impact this critical shell-to-ring transition. In previous work, we used an agent-based model to examine the geometrical arrangement of growing actin filaments in the presence of VASP, which was represented as a tetrameric actin-binding protein. When placed inside a spherical container to mimic the condensate geometry, VASP promoted the assembly of actin filaments into ring-like bundles.^18^ These simulations revealed that VASP can form bundles of actin in kinetic regimes characterized by a slightly higher rate of VASP-actin binding than unbinding.^18^ However, when the rate of unbinding was higher than the rate of binding, the actin filaments failed to form a ring, remaining in a shell-like arrangement. Here we adapted this model to investigate whether a bivalent actin-binding protein, similar to Lamellipodin, could also drive the formation of actin rings in the condensate environment, where the protein is locally concentrated in the presence of actin filaments (**Fig. S1**). Our simulations revealed a kinetic regime in which bivalent actin-binding proteins drive actin filaments to undergo a shell-to-ring transition **(Fig. 4I-K)**. To monitor this transition in our simulations, we quantified two metrics.^18^ First, we observed that the presence of bivalent actin crosslinkers causes a reduction in the fraction of the inner condensate surface occupied by filaments, as would be expected for a shell-to-ring transition **(Fig. 4J).** Second, we found that as rings formed, the fraction of bivalent crosslinkers bound to two filaments increased **(Fig. 4K)**. We note that our modeling approaches have some limitations because we model the condensate as a spherical container with rigid boundaries. However, as we showed previously,^18^ this assumption does not impact our conclusions about the assembly of shell and ring morphologies, as these structures form during the period when the condensate is still approximately spherical. These results collectively suggest that within the condensate environment, where the concentration of proteins is high, a dimeric actin binder can sufficiently cross-link actin filaments to drive the formation of ring-like bundles, which, in experiments, eventually deform and elongate condensates to form linear actin bundles. Further, to mimic condensates consisting of VASP and Lamellipodin **(Fig. 3)**, we simulated mixtures of tetravalent and bivalent crosslinkers, which also resulted in a shell-to-ring transition, as expected (**Fig. S2**, see also **Supplementary Movie M1**). Our simulations reveal that the ratio of tetravalent to bivalent crosslinkers can tune the balance between shells and rings (**Fig. S3**).

### Dynamic dimerization of crosslinkers is sufficient to bundle actin filaments within condensates

Having observed in both experiments and simulations that mini-Lpd is sufficient to drive actin ring formation and bundling, we wondered to what extent these behaviors depend on the dimeric nature of mini-Lpd. Protein condensates are known to promote multivalent interactions among proteins,^39,51,52^ yet stable multimerization of the constituent proteins is not required for condensate formation. On the contrary, some of the best studied examples of condensate forming proteins are monomeric under dilute conditions.^10,53^ Therefore, we asked whether the inherent multi-valency of the condensate environment might be sufficient to drive actin bundling. To probe this point, we simulated monomeric actin-binding proteins that have an affinity for one another and for actin, such that they form dynamically reversible dimers, as may occur when they bind to closely spaced actin filaments **(Fig. 5A)**. This dynamic dimerization model has two sets of binding affinities – one for actin binding to the monomer and another for dimerization of the monomer, **Fig. 5A**. We investigated the role of dimerization in a regime where actin is already known to form rings with stable dimeric crosslinkers (**Fig. 4H-J**, k_bind_ = 10.0 s^-1^, k_unbind_ = 1.0 s^-1^). Our simulations reveal that even with dynamically dimerizing monomers, actin filaments can form bundles given a sufficiently long dimer residence time (**Fig 5B**). In the case of dynamic dimerization, the formation of rings depends on the balance between dimer formation and dimer splitting. When the rate of dimer formation is higher than dimer splitting, the fraction of the condensate inner surface area covered by actin is low, corresponding to a ring state (**Fig. 5C** top), similar to our observations for permanent dimers (**Fig. 4G)**. In contrast, when the dimer splitting rate is higher than the dimer formation rate, we observe shells (**Fig. 5C** bottom). Thus, ring formation is favored when dimers have a higher residence time. We next examined the distribution of monomers and dimers bound to actin in these simulations. The actin binding protein can exist in four subpopulations: free monomers, free dimers, actin-bound monomers, and actin-bound dimers. We found that in cases where actin shells were predominant, the fraction of monomers bound to actin was higher, while the fraction of dimers bound to actin was negligible. In contrast, conditions that supported the assembly of ring-like actin bundles had a higher fraction of actin bound dimers (**Fig. 5D**). Here, we chose a propensity for dimer formation as the simplest representation of protein clustering in the condensate environment. The resulting trend of increasing ring formation with increasing dimer affinity would be expected to increase if higher order multimerization, as is likely present in the condensate environment, were included in the simulations.

**Figure 5:**
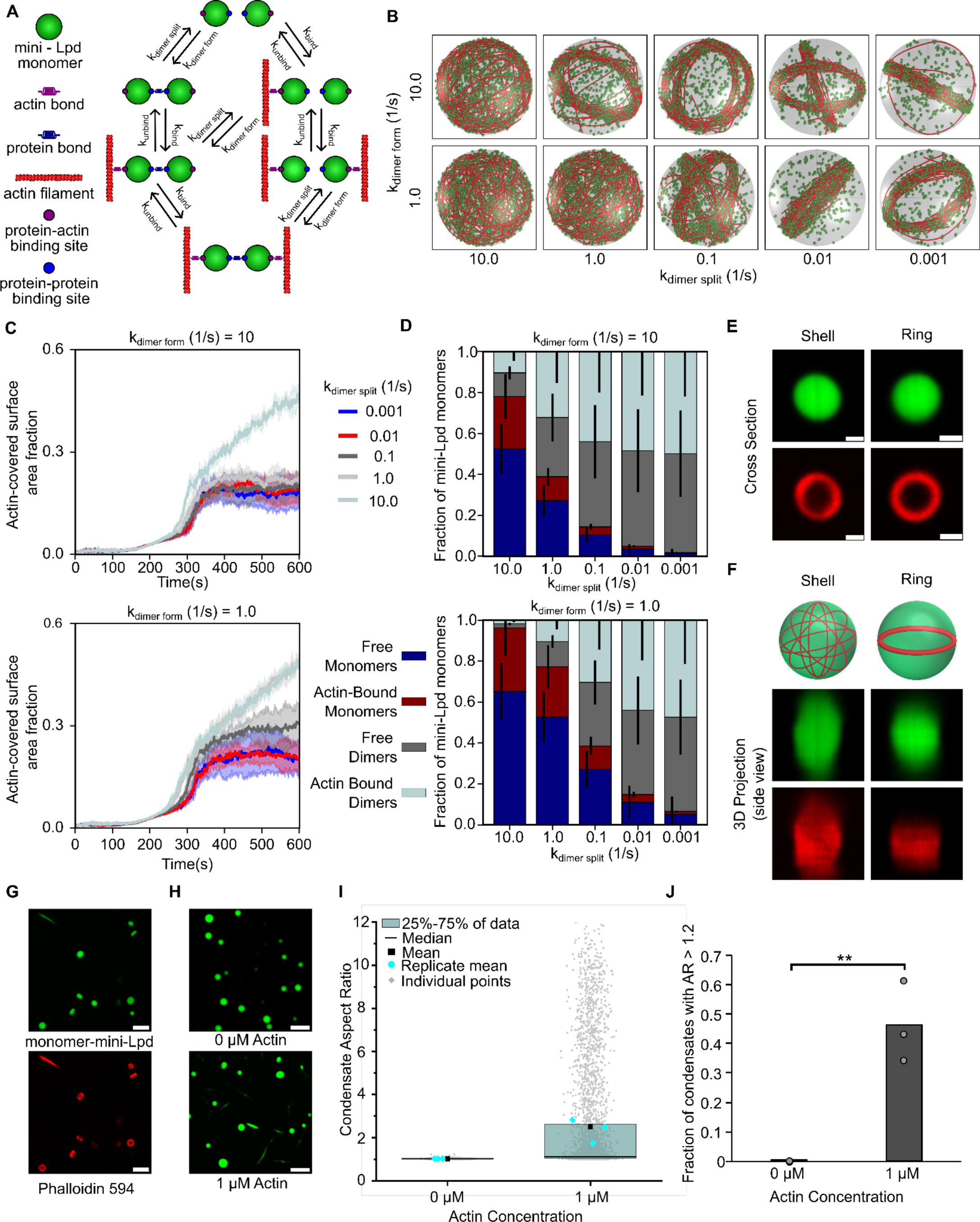
Dynamic protein dimers bundle actin filaments. **A)** Schematic showing the different possible binding interactions between the monomeric actin binding protein and actin, including both protein-protein interactions (dynamic dimerization) and protein-actin interactions. **B)** Simulations show that monovalent actin binding can lead to the formation of ring structures when monomers are allowed to interact, bind to form dimers, and function as transient bivalent crosslinkers. Representative final snapshots (t = 600 s) from simulations at various dimer formation and splitting rates within spherical condensates (R = 1 μm) containing 30 actin filaments (red) and 2000 monomers (green spheres) which can form a maximum of 1000 dimers. Please refer to the Supplementary Methods section for a detailed description of the model. The dimer formation rates of the monomers are varied along each column, and dimer splitting rates are varied along each row. Actin-binding kinetics were chosen from previous simulations (Fig. 4I) to correspond to ring-forming conditions for bivalent crosslinkers. The polymerization rate at the plus (+) end is constant at 0.0103 μm/s, and neither end undergoes depolymerization. Also see **Supplementary Movie M4**. **C)** Time series showing the mean (solid line) and standard deviation (shaded area) of simulated condensate surface that is covered with actin. The k_dimer form_ value is shown on top of each subpanel while time series are colored by k_dimer split_ values. Data used: 5 replicates. Please refer to supplementary methods for a detailed description. **D)** Stacked bar graph showing the distribution of condensate protein at different states namely, free monomers, free dimers, actin-bound monomers, and actin-bound dimers. Error bars show standard deviation. Data used: 5 replicates, data from last 30 snapshots. **E)** Representative 2D confocal cross sections of independent monomer mini-Lpd (green) condensates containing peripheral actin, shown with phalloidin staining (red), in both a shell of actin (top) and a ring of actin (bottom). Scale bars 1 μm. **F)** 3D reconstructions of the same condensates shown in **E** demonstrating a shell of actin (top) and a ring of actin (bottom). **G)** Phalloidin-iFluor-594 (red) staining of monomer-mini-Lpd condensates (green) with 5 μM monomeric G-actin (unlabeled) added displaying polymerized actin within the protein condensates. Scale bars 5 μm. **H)** The addition of actin to monomer-mini-Lpd also results in the deformation of protein condensates formed from 5 μM monomer-mini-Lpd. Scale bars 5 μm. **I)** Distribution of condensate aspect ratios across the conditions in **H**, with at least 1000 condensates analyzed for each condition. For the 1 μM actin condition values for aspect ratios above 10, corresponding to 2.97% of the data, are not displayed to better visualize distributions for all conditions. The mean is the mean of all data points in each condition, while the replicate mean is the mean of each of the individual replicates for each condition. **J)** Quantification of the fraction of high-aspect-ratio protein condensates, defined as condensates with aspect ratios > 1.2, across the conditions in **H**. Data are mean across three independent experiments with at least 1000 condensates analyzed per condition. Overlayed gray circles denote the means of each replicate. Two asterisks denote p<.01 using an unpaired, two-tailed t-test on the means of the replicates N=3. All experiments were performed in a buffer containing 20 mM Tris (pH 7.4), 50 mM NaCl, 5 mM TCEP, and 3% (w/v) PEG 8000.

To test the prediction that a stable multimer is not required for actin bundling in protein condensates, we examined a monomeric version of mini-Lpd, which we will refer to as monomer-mini-Lpd. This protein lacked the dimerizing leucine zipper domain. In line with the model predictions, condensates formed from monomer-mini-Lpd behaved similarly to those of the dimeric mini-Lpd upon exposure to actin. Upon the addition of monomeric actin, actin filaments began to polymerize inside the condensates and partitioned to the inner condensate surface where they assembled into shells and ring-like bundles **(Fig. 5E,F).** As actin continued to polymerize inside the condensates, confirmed through phalloidin staining **(Fig. 5G)**, the condensates were progressively deformed **(Fig. 5H)** and had high aspect ratios **(Fig. 5I,J)**, similar to those formed upon the addition of actin to condensates of VASP,^13^ Vasp/mini-Lpd (**Fig. 3A**), and mini-Lpd (**Fig. 4A**). The ability of condensates composed of proteins that lack any known polymerase activity, mini-Lpd and monomer-mini-Lpd, to polymerize and bundle actin filaments led us to ask how protein condensates might facilitate the assembly of actin filaments. Multivalent binding to actin filaments is thought to be a key functional requirement for the two major classes of known polymerases, formins and members of the ENA/VASP family.^7,32,49^ Formins are native dimers, while ENA/VASP proteins are native tetramers.^7,14^ Both polymerases function by binding simultaneously to actin filaments and monomeric actin, resulting in the addition of monomers to the barbed ends of growing filaments. Given the essential role of multivalent binding in actin polymerization, we wondered whether protein condensates, which inherently promote multivalent protein contacts,^8,9,39^ might have an inherent capacity to promote actin polymerization.

### Adding an actin-binding domain to a condensate-forming protein confers the ability to polymerize and bundle actin filaments

To probe the contribution of the condensate environment to actin assembly, we sought to identify the minimum requirements for actin polymerization and bundling by protein condensates. In our previous work with VASP, condensates composed of a VASP mutant lacking the G-actin binding site retained the ability to polymerize and bundle actin filaments. In contrast, condensates that lacked the F-actin binding site or lacked both F-actin and G-actin binding sides failed to polymerize actin or deform upon actin addition.^13^ Therefore, the minimum requirement for actin polymerization and bundling by protein condensates could be the ability to bind filamentous actin. Given that condensates are composed of a dense network of interconnected proteins, actin filaments that reside inside a condensate are likely to be in contact simultaneously with many copies of the condensate-forming protein. For this reason, the condensate-forming protein need not be an inherent multimer to drive actin polymerization, as we observed with condensates of monomer-mini-Lpd, **Fig. 5**. Therefore, the binding of multiple condensate proteins to a growing actin filament could function similarly to the multivalent binding of filaments by multimeric actin polymerases, effectively meeting the key requirement for actin polymerization.

To investigate this possibility, we examined the interaction of actin with a condensate-forming protein that has no known ability to bind or polymerize actin and then conferred actin-binding ability upon it through the addition of an actin-binding domain. The protein we selected for this experiment was Eps15, a protein involved in clathrin-mediated endocytosis, which has no known interaction with actin.^54–56^ Eps15 has an N-terminal region consisting of three structured EH domains, a central coiled-coil domain, through which the protein forms native dimers, and an intrinsically disordered C-terminal domain.^37^ Binding interactions between the N and C-terminal domains drive Eps15 to form protein condensates in vitro **(Fig. 6A)**.^35^ As expected, the addition of 3 μM monomeric actin to Eps15 condensates did not result in actin polymerization, as no filaments were observed upon phalloidin staining and the condensates did not deform **(Fig. 6B).** We then fused a filamentous actin-binding motif, the 17 amino acid Lifeact peptide, to the C-terminus of Eps15 to create Eps15-Lifeact **(Fig. 6C)**. Lifeact, which binds to actin filaments at the interface between two monomers, is commonly used in conjunction with fluorophores to visualize the filamentous actin cytoskeleton.^57–59^ It does not alter the bulk polymerization rate of growing actin filaments,^57^ though it can increase the initial rate of barbed end elongation, likely by stabilizing nascent filaments.^60^ Mixing wild-type Eps15 and Eps15-Lifeact at a 1:1 ratio (15 μM total protein) in the presence of 3% PEG (w/v) led to the co-condensation of the two proteins. Condensates were formed using this 1:1 ratio to minimize any potential effect of Lifeact on Eps15 phase separation. The partitioning of wild-type Eps15 into these condensates was similar to that of condensates consisting purely of wild-type Eps15, suggesting that Lifeact addition had little effect on Eps15 phase separation (**Fig. 6D).** When monomeric actin was added to the resulting condensates, they deformed into high aspect ratio structures. Actin polymerization within these condensates was confirmed by phalloidin staining, which revealed their progressive deformation from spherical to ellipsoid to rod-like morphologies **(Fig. 6F,G).** Prior to these morphological transitions, actin shells and ring-like bundles formed within spherical condensates **(Fig. 6H,I),** as described above for condensates of mini-Lpd **(Fig. 4G,H)**, and monomer-mini-Lpd **(Fig. 5E,F)**, and previously for condensates of VASP.^13^ Quantification of condensate morphologies confirmed that increasing concentrations of monomeric actin drove a substantial increase in the aspect ratios of condensates (**Fig. 6I,J)**. These results demonstrate that it is possible to confer actin polymerization and bundling activity upon an arbitrarily chosen condensate-forming protein, simply by adding a filamentous actin binding domain to it.

**Figure 6.**
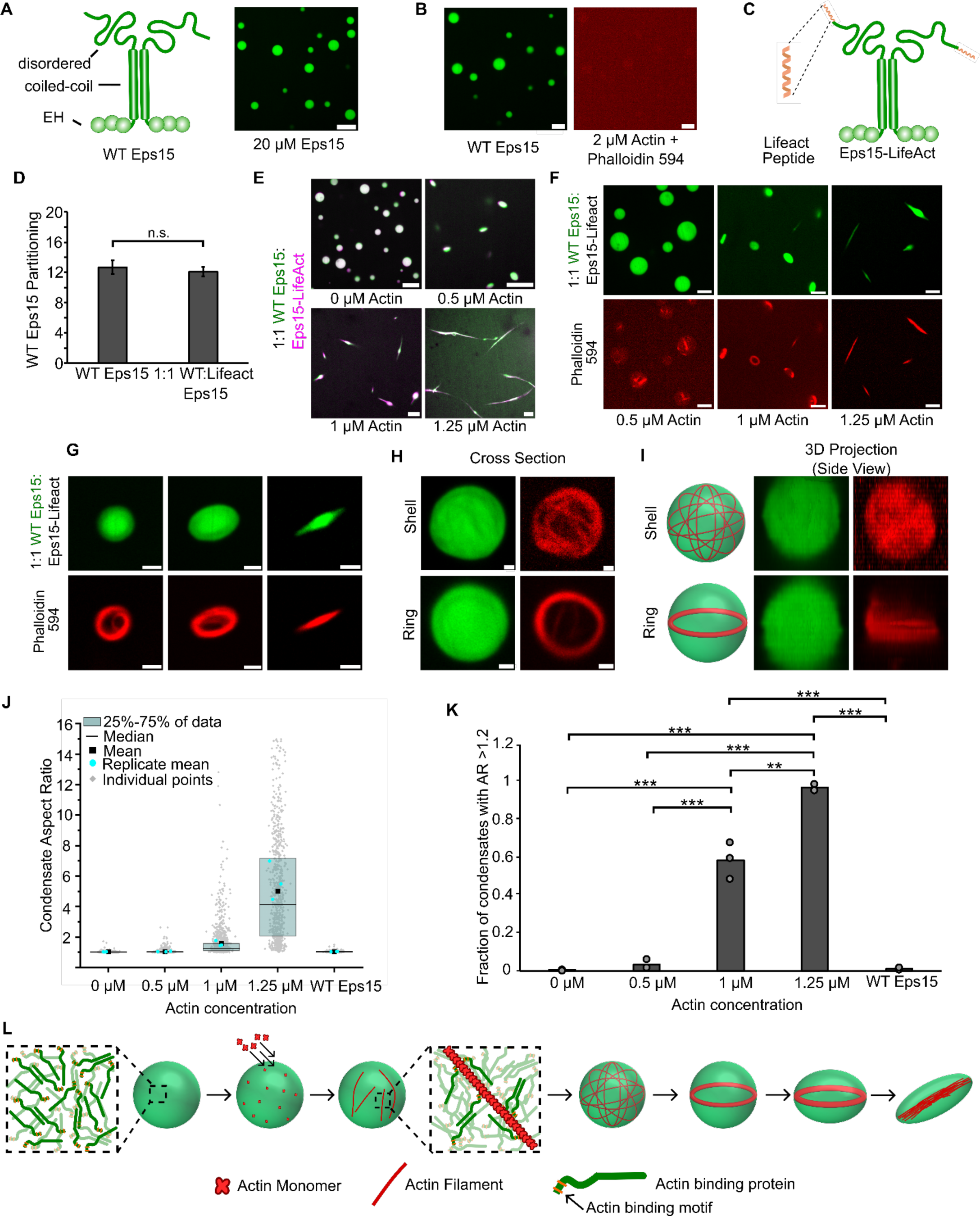
Adding an actin-binding domain to an arbitrary condensate-forming protein is sufficient to confer the ability to polymerize and bundle actin filaments. **A)** Left: Schematic depicting wild type Eps15 and its major domains. Right: 20 μM wild type Eps15 forms condensates in solution with 3% (w/v) PEG. Scale bar 5 μm. **B)** Wild-type Eps15 (green) condensates do not polymerize or bundle actin as indicated by the lack of condensate deformation and a lack of phalloidin-stained filaments. Scale bars 5 μm. **C)** Schematic depicting fusion of the Lifeact peptide to the C-terminus of Eps15. **D)** Partitioning of wild type Eps15 in condensates formed from only wild type Eps15 (self-partitioning) and condensates formed with 1:1 wild type Eps15 and Eps15-Lifeact. Six fields of view were analyzed for each condition with more than 6 condensates analyzed for partitioning per field of view. Error bars indicate standard deviation. **E)** Representative images of condensate deformation and actin polymerization upon the addition of increasing concentrations of monomeric actin to condensates formed from wild-type Eps15 and Eps15-Lifeact. Scale bars 5 μm. **F)** Representative images of phalloidin staining of condensates consisting of wild-type Eps15 and Eps15-Lifeact following actin addition. Scale bars 5 μm. **G)** Representative confocal cross-section images of the progressive, actin-driven deformation of condensates consisting of wild-type Eps15 and Eps15-Lifeact. Scale bars 2 μm. **H)** Representative 2D confocal images of independent 1:1 WT Eps15:Eps15-Lifeact condensates (green) containing peripheral actin, shown with phalloidin staining (red), in both a shell of actin (top) and a ring of actin (bottom). Scale bars 1μm. **I)** 3D reconstructions of the same condensates shown in **H** demonstrating a shell of actin (top) and a ring of actin (bottom). **J)** Distribution of condensate aspect ratios across the conditions in **E**, with at least 700 condensates analyzed for each condition. For the 1.25 μM actin condition, values for aspect ratios higher than 15, corresponding to 3.5% of the data, are not displayed to better visualize distributions for all conditions. The mean is the mean of all data points in each condition, while the replicate mean is the mean of each of the individual replicates for each condition. **K)** Quantification of the fraction of elongated protein condensates, defined as condensates with aspect ratios > 1.2, across the conditions in **E**. Data are mean +/- standard deviation across three independent experiments with at least 700 condensates analyzed in total per condition. Overlayed gray circles denote the means of each replicate. Two asterisks denote p<.01, and three asterisks denote p<.001 using an unpaired, two-tailed t-test on the means of the replicates N=3. **L)** Cartoon depicting the proposed mechanism of condensate-driven actin polymerization and bundling by condensates of actin-binding proteins. All experiments were performed in a buffer containing 20 mM Tris (pH 7.4), 150 mM NaCl, 5 mM TCEP, and 3% (w/v) PEG 8000.

## Discussion

Taken together, our findings suggest that the ability to bind filamentous actin is sufficient to drive actin polymerization and bundling by protein condensates through the mechanism outlined in **Fig. 6K**. Condensates formed from actin-binding proteins are capable of locally concentrating actin. As the actin concentration increases inside condensates, the polymerization of actin filaments results from multivalent interactions between filament nuclei and actin-binding motifs within condensate proteins. While the precise mechanism of filament nucleation was not revealed in our studies, concentrating actin above a threshold concentration is known to promote filament nucleation and polymerization.^3,61,62^ Nascent filaments are likely stabilized by multivalent contacts with condensate proteins that contain filamentous actin binding sites.^7,60^ As actin filaments grow within condensates, they spontaneously partition to the edges of condensates to minimize filament curvature energy. This phenomenon has been reported whenever filaments grow inside spherical containers with diameters below the persistence length of actin, 10-20 μm.^13,18,63–65^ This partitioning results in the assembly of actin shells at the inner surfaces of the condensate, which rearrange to form ring-like actin bundles. As more filaments join these structures, their rigidity eventually overcomes the surface tension of the condensate, permitting the filaments to straighten and thereby deforming the condensate into a rod-like structure filled with a bundle of parallel actin filaments.^13^ Here we have used a combination of experiments and agent-based computational modeling to illustrate that this process does not require stable multimerization of the condensate-forming protein. Provided they bind to actin filaments, transient interactions between condensate-forming proteins appear sufficient to drive actin bundling.

By illustrating that these steps can occur in the absence of proteins with inherent polymerase activity, our findings suggest that the set of proteins involved in the assembly and organization of actin filaments may be substantially larger than previously thought. Actin has a large interactome consisting of more than 100 proteins,^66^ only a small fraction of which are known to polymerize actin filaments. The existing understanding of actin polymerization focuses on specialized polymerase proteins such as formins and members of the ENA/VASP family.^7,14,33^ While these proteins have distinct structures and mechanisms, they share a common ability to make flexible, multi-valent contacts with actin filaments, which is essential to their function.^7,13,14,32^ Our studies suggest that the condensate environment has an inherent capacity to promote these flexible, multi-valent contacts. As a result, the individual proteins that make up condensates need not have dedicated polymerase activity to influence actin polymerization, but simply the ability to bind actin.

Additionally, many actin-interacting proteins contain both proline-rich motifs and proline-binding domains, such as SH3, EVH1, or WW domains.^14,39,67,68^ Interactions between these domains and sequences often lead to the assembly of multivalent protein networks, which are the building blocks of liquid-like condensates.^11,39,69^ In line with this reasoning, recent work in several labs has shown that many actin-interacting proteins participate in condensate networks or form condensates themselves and that many of these condensates facilitate actin polymerization.^11,13,23,38–42^ In this context, our results suggest that the ability to assemble and organize the actin cytoskeleton is an emergent property of liquid-like, multivalent protein networks, rather than the sole responsibility of specialized polymerase proteins.

## Methods

### Reagents

Tris base, NaCl, Tris(2-carboxyethyl)phosphine (TCEP), poly-L-lysine, and Atto 594 maleimide were purchased from Sigma-Aldrich. Alexa Fluor 647 C2 maleimide was purchased from Thermo Fischer Scientific. Phalloidin-iFLuor 594 was purchased from Abcam. Amine-reactive PEG (mPEG– succinimidyl valerate, MW 5,000) was purchased from Laysan Bio. Rabbit muscle actin was purchased from Cytoskeleton.

### Plasmids

A pET vector encoding the ‘cysteine light’ variant of human VASP (pET-6xHis-TEV-KCK-VASP(CCC-SSA)) was a gift from Scott Hansen. All VASP mutants were generated using this plasmid as a template, as previously described.^13^ Briefly, monomeric VASP (mVASP) was generated using site-directed mutagenesis to introduce a stop codon after amino acid 339 to truncate VASP and remove its tetramerization domain. ΔEVH1 VASP was generated using PCR to delete the EVH1 domain (amino acids 1-113) before recircularization through restriction enzyme digestion and ligation.

The vector encoding mini-Lpd (his-Z-EGFP-LZ-Lpd(aa850-1250)) and monomeric-mini-Lpd (his-EGFP-Lpd(aa850-1250)) were gifts from Scott Hansen.

pET28a 6×His-Eps15 (FL), encoding H. sapiens Eps15, was a gift from Tomas Kirchhausen. Lifeact addition to Eps15 FL was done through site-directed mutagenesis. PCR using oligonucleotides 5’ TTCGAGAGCATCAGCAAGGAAGAGTGAGGATCCGAATTCGAGCTCC 3’ and 5’ CTTCTTGATCAGGTCGGCCACGCCCATTGCTTCTGATATCTCAGATTTGCTGAGTG 3’ was done to introduce the Lifeact sequence to the c-terminal end of Eps15. PCR was followed by the addition of the PCR product to a KLD enzyme mix (NEB) for template removal, phosphorylation, and ligation into a re-circulated plasmid.

### Protein Purification

The mini-Lpd (his-Z-EGFP-LZ-Lpd(aa850-1250)) and monomeric-mini-Lpd (his-EGFP-Lpd(aa850-1250)) were transformed into BL21 (NEB, cat. No. C2527H) and grown at 30 °C to an OD of 0.8. The bacteria were then cooled to 12°C and induced for 24 hours with 1mM IPTG. The rest of the protocol was performed at 4 °C. Cells were pelleted from a 2L culture by centrifugation at 4,785g (5,000 rpm in Beckman JLA-8.100) for 20 min. Pellets were resuspended in 100mL of lysis buffer (50mM sodium phosphate pH 8.0, 300mM NaCl, 10mM imidazole, 0.5mM TCEP, 0.2% Triton X100, 10% glycerol, 1mM PMSF, and EDTA free protease inhibitor tablets (1 tablet per 50mL) (Roche cat# 05056489001)) followed by sonication on ice for 4×2000J with amplitude at 10 (Sonicator Qsonica LLC, Q700). The lysate was clarified by centrifugation at 48,384g (20,000 rpm in Beckman JA25.50) for 30 min at 4 °C before being applied to a 10mL bed volume Nickel nitrilotriacetic acid (Ni-NTA) agarose (Qiagen, cat. no. 30230) column, and washed with 10 column volumes (CVs) of lysis buffer to which imidazole had been added to a final concentration of 20mM. The column was then washed with 5 column volumes of lysis buffer containing 20 mM Imidazole but lacking Triton-X100 and protease inhibitor tablets. The protein was eluted with elution buffer (50 mM Tris-HCl, pH 7.5, 300mM NaCl, 10% glycerol, 400mM imidazole, 1mM TECP, 1mM PMSF, and EDTA-free protease inhibitor tablets (1 tablet per 50mL). The protein was concentrated using Amicon Ultra-15, 30K MWCO (Millipore: Cat#UFC903024) to 5 mL, and clarified by ultracentrifugation for 5min at 68,000 x g (40,000 rpm with Beckman optimal MAX-E Ultracentrifuge and TLA100.3 rotor). The protein was further purified by size exclusion chromatography with Superose 6, and ion exchange chromatography with SP Sepharose Fast Flow (GE Healthcare, Cat#17-0729-01), and stored as liquid nitrogen pellets at -80°C.

The pET-His-KCK-VASP(CCC-SSA) plasmid was transformed into Escherichia coli BL21(DE3) competent cells (NEB, cat. no. C2527). Cells were grown at 30 °C to an optical density (OD) of 0.8. Protein expression was performed as described previously with some alteration.^32^ Expression of VASP was induced with 0.5 mM isopropylthiogalactoside (IPTG), and cells were shaken at 200 rpm at 12 °C for 24 h. The rest of the protocol was carried out at 4 °C. Cells were pelleted from 2 L cultures by centrifugation at 4,785g (5,000 rpm in Beckman JLA-8.100) for 20 min. Cells were resuspended in 100 mL lysis buffer (50 mM sodium phosphate pH 8.0, 300 mM NaCl, 5% glycerol, 0.5 mM TCEP, 10 mM imidazole, 1 mM phenylmethyl sulphonyl fluoride (PMSF)) plus EDTA-free protease inhibitor tablets (1 tablet per 50 mL, Roche, cat. no. 05056489001), 0.5% Triton-X100, followed by homogenization with a dounce homogenizer and sonication (4 × 2,000 J). The lysate was clarified by ultracentrifugation at 125,171g (40,000 rpm in Beckman Ti45) for 30 min. The clarified lysate was then applied to a 10 mL bed volume Nickel nitrilotriacetic acid (Ni-NTA) agarose (Qiagen, cat. no. 30230) column, washed with 10 column volumes of lysis buffer plus EDTA-free protease inhibitor tablets (1 tablet per 50 mL), 20 mM imidazole, 0.2% Triton X-100, followed by washing with 5×CV of lysis buffer plus 20 mM imidazole. The protein was eluted with elution buffer (50 mM Tris, pH 8.0, 300 mM NaCl, 5% glycerol, 250 mM imidazole, 0.5 mM TECP, EDTA-free protease inhibitor tablets (1 tablet per 50 mL)). The protein was further purified by size exclusion chromatography with Superose 6 resin. The resulting purified KCK-VASP was eluted in storage buffer (25 mM HEPES pH 7.5, 200 mM NaCl, 5% glycerol, 1 mM EDTA, 5 mM DTT). Single-use aliquots were flash-frozen using liquid nitrogen and stored at −80 °C until the day of an experiment. The his-tagged VASP mutants were purified using the same protocol as above as indicated or with the following modifications: His-KCK-VASPΔTet: no modifications. GST-KCK-VASPΔEVH1 was purified using the same protocol as above but with the following buffer modifications: The lysis buffer was 20 mM Tris pH 8.5, 350 mM NaCl, 5% glycerol, 5 mM EDTA, 5 mM DTT, and 1 mM PMSF. The storage buffer was 25 mM HEPES, pH 7.5, 200 mM NaCl, 5% glycerol, 1 mM EDTA, and 5 mM DTT

Full-length Eps15 and Eps15-Lifeact were expressed as N-terminal 6x-His-tagged constructs in BL21(DE3) *E. Coli* cells. Cells were grown in 2xYT medium for 3-4 hours at 30 °C to an optical density at 600 nm of 0.6-0.9, cooled for 1 hour, and then protein expression was induced with 1 mM IPTG at 12°C for 20-30 hours. Cells were collected, and bacteria were lysed in a lysis buffer using homogenization and probe sonication. Lysis buffer was 50 mM Tris-HCl, pH 8.0, 300 mM NaCl, 5 mM imidazole, 10 mM β-mercaptoethanol or 5 mM TCEP, 1 mM PMSF, 0.2% Triton X-100 and 1x Roche or Pierce complete EDTA-free protease inhibitor cocktail tablet per 50 mL buffer. Proteins were incubated with Ni-NTA Agarose (Qiagen 30230) resin, followed by extensive washing with 10 column volumes, then eluted from the resin in 50 mM Tris-HCl, pH 8.0, 300 mM NaCl, 200 mM imidazole, 10 mM β-mercaptoethanol or 5 mM TCEP, 1 mM PMSF, and 1× Roche or Pierce complete EDTA-free protease inhibitor cocktail tablet. The protein was then further purified by gel filtration chromatography using a Superose 6 column equilibrated with 20 mM Tris-HCl, pH 8.0, 150 mM NaCl, 1 mM EDTA, and 5 mM DTT. Purified proteins were concentrated using Amicon Ultra-15 Ultracell-30K centrifugal filter units (Millipore–Sigma), then centrifuged at 100,000 rpm at 4 °C for 10min using a Beckman TLA-120.2 rotor to remove aggregates, and stored either in small aliquots or as liquid nitrogen pellets at −80 °C.

### Protein labeling

The VASP used in these studies is a previously published ‘cysteine light’ mutant that replaced the three endogenous cysteines with two serines and an alanine. A single cysteine was then introduced at the N-terminus of the protein to allow selective labeling with maleimide dyes. This mutant was found to function in an indistinguishable manner from the wild-type proteins.^14^ Thus, VASP and its mutants were labeled at the N-terminal cysteine using maleimide-conjugated dyes. VASP was buffer exchanged into 20 mM Tris (pH 7.4) 150 mM NaCl buffer to remove DTT from the storage buffer and then incubated with a three-fold molar excess of dye for two hours at room temperature. Free dye was then removed by applying the labeling reaction to a Zeba Dye and Biotin removal size exclusion column (Thermo Fischer Scientific) equilibrated with buffer containing 20 mM Tris, 150 mM NaCl, and 5 mM TCEP pH 7.4.

Monomeric actin was labeled using maleimide-conjugated dyes. Dyes were incubated with G-actin at a 2-fold molar excess for 2 hours at room temperature before being separated from the labeled protein by applying the labeling reaction to a spin column packed with Sephadex G-50 Fine DNA Grade (GE Healthcare GE17-0573-01) hydrated with A buffer (5 mM Tris-HCL (pH 8), 0.2 mM ATP and 0.5 mM DTT pH 8). The labeled protein was then centrifuged at 100,000 x G for 10 min at 4 degrees Celsius to remove aggregates before being flash-frozen in single-use aliquots.

Eps15 and Eps15-Lifeact were labeled using amine-reactive NHS-ester dyes at a 3-fold molar excess of dye before free dye was removed by applying the labeling reaction to a Zeba Dye and Biotin removal size exclusion column (Thermo Fischer Scientific) equilibrated with buffer containing 20 mM Tris (pH 7.4), 150 mM NaCl, and 5 mM TCEP.

### Protein condensate formation and actin polymerization

Condensates composed of either VASP or mini-Lpd/monomer mini-Lpd alone were formed by mixing the given concentration of protein (see text) with 3% (w/v) PEG 8000 in 20 mM Tris pH 7.4, 5 mM TCEP, and the given concentration of NaCl (50 mM for mini-Lpd/monomer mini-Lpd and 150 mM for mini-Lpd + VASP, VASP, or Eps15). PEG was added last to induce condensate formation after the protein was evenly dispersed in the solution. For condensates consisting of both mini-Lpd and VASP, formed in the absence of PEG, the only difference was that PEG 8000 was not added to the mix, however, all other buffer conditions were as stated above. All protein concentrations listed are the monomeric concentrations.

For actin polymerization assays within condensates consisting of VASP, mini-Lpd, or Eps15, condensates were formed and then G-actin was added to the condensate solution and allowed to polymerize for 15 minutes before imaging. For phalloidin-actin assays, unlabelled G-actin was added to pre-formed protein condensates and allowed to polymerize for 10 min. Phalloidin-iFluor594 (Abcam) was then added to stain filamentous actin for 10 min before imaging. For assays that included Latrunculin, 5 μM Latrunculin A (Cayman Chemical Item No.10010630) was added to the pre-formed protein condensates and mixed gently before actin addition.

For FRAP experiments, condensates formed from the various proteins were observed in solution at the conditions given in the text. A region within the condensates was bleached and consecutive images were taken every three seconds to monitor fluorescence recovery over time.

### Microscopy

Samples were prepared for microscopy in wells formed from 1.6 mm thick silicone gaskets (Grace Biolabs) on Hellmanex II (Hellma) cleaned, no. 1.5 glass coverslips (VWR). Coverslips were passivated using poly-L-lysine conjugated PEG chains (PLL-PEG). To prevent evaporation during imaging, an additional small coverslip was placed on top of the gasket to seal the well. Fluorescence microscopy was done using an Olympus SpinSR10 spinning disk confocal microscope with a Hamamatsu Orca Flash 4.0V3 Scientific CMOS camera. FRAP was done using the Olympus FRAP unit 405 nm laser.

PLL-PEG was prepared as described previously with minor alterations. Briefly, amine-reactive mPEG succinimidyl valerate was conjugated to poly-L-lysine at a molar ratio of 1:5 PEG to PLL. The conjugation reaction takes place in 50mM sodium tetraborate solution pH 8.5 and is allowed to react overnight at room temperature while continuously stirring. The final product is then buffer exchanged to PBS pH 7.4 using 7000 MWCO Zeba spin desalting columns (Thermo Fisher) and stored at 4 °C.

### Image Analysis

Image J was used to quantify the distribution of condensate characteristics. Specifically, condensates were selected using thresholding in the brightest channel and shape descriptors (i.e. diameter, aspect ratio, etc.), and protein fluorescent intensities were measured using the built-in analyze particles function.

FRAP data were analyzed using ImageJ where fluorescence recovery over time was measured and then normalized to the maximum pre-bleach intensity. Recovery was measured for condensates of similar diameters and photobleached region size.

Partitioning data was calculated using the average intensities of the condensed protein phase and the bulk solution, with partitioning defined as the ratio of the intensity inside the condensate to outside the condensate. Images were cropped so that only condensates from the middle ninth of the field of view were analyzed to avoid any error from potential non-uniform illumination across the imaging field.

## Acknowledgments

This research was supported by the National Science Foundation through the Center for Dynamics and Control of Materials: an NSF MRSEC under Cooperative Agreement No. DMR-2308817. Additionally, this research was supported by grants from the NIH to J.C.S (R35GM139531), P. R. (R01GM132106) and by the NSF through a Modulus Grant MCB 2327244 to P.R and J.C.S.

## Contributions

C.W, K.G, A.C., D.M, P.R., and J.C.S. designed experiments. C.W., K.G., A.C., D.M, P.R., and J.C.S. wrote and edited the manuscript. C.W, K.G, A.C., D.M, A.T., P.R., and J.C.S. performed experiments and analyzed data. All authors consulted on manuscript preparation and editing.

## Corresponding authors

Correspondence to Padmini Rangamani or Jeanne C. Stachowiak.

## Notes

### Competing Interest Statement

The authors have declared no competing interest.

### Summary of Updates

We have updated this manuscript to include agent-based computational modeling, led by the Rangamani group. For this reason, two new authors, Chandrasekaran and Mansour have been added.

## References

1. Fletcher, D. A. & Mullins, R. D. Cell mechanics and the cytoskeleton. Nature 463, 485–492 (2010).

2. Hinze, C. & Boucrot, E. Local actin polymerization during endocytic carrier formation. Biochem. Soc. Trans. 46, 565–576 (2018).

3. Pollard, T. D. & Cooper, J. A. Actin, a Central Player in Cell Shape and Movement. Science 326, 1208– 1212 (2009).

4. Vasioukhin, V., Bauer, C., Yin, M. & Fuchs, E. Directed Actin Polymerization Is the Driving Force for Epithelial Cell–Cell Adhesion. Cell 100, 209–219 (2000).

5. Campellone, K. G. & Welch, M. D. A nucleator arms race: cellular control of actin assembly. Nat. Rev. Mol. Cell Biol. 11, 237–251 (2010).

6. Dominguez, R. Actin filament nucleation and elongation factors – structure–function relationships. Crit. Rev. Biochem. Mol. Biol. 44, 351–366 (2009).

7. Chesarone, M. A., DuPage, A. G. & Goode, B. L. Unleashing formins to remodel the actin and microtubule cytoskeletons. Nat. Rev. Mol. Cell Biol. 11, 62–74 (2010).

8. Alberti, S., Gladfelter, A. & Mittag, T. Considerations and Challenges in Studying Liquid-Liquid Phase Separation and Biomolecular Condensates. Cell 176, 419–434 (2019).

9. Boeynaems, S. et al. Protein Phase Separation: A New Phase in Cell Biology. Trends Cell Biol. 28, 420–435 (2018).

10. Elbaum-Garfinkle, S. et al. The disordered P granule protein LAF-1 drives phase separation into droplets with tunable viscosity and dynamics. Comput. Biol. 6.

11. Su, X. et al. Phase separation of signaling molecules promotes T cell receptor signal transduction. Science 352, 595–599 (2016).

12. Woodruff, J. B. et al. The Centrosome Is a Selective Condensate that Nucleates Microtubules by Concentrating Tubulin. Cell 169, 1066–1077.e10 (2017).

13. Graham, K. et al. Liquid-like VASP condensates drive actin polymerization and dynamic bundling. Nat. Phys. 19, 574–585 (2023).

14. Krause, M., Dent, E. W., Bear, J. E., Loureiro, J. J. & Gertler, F. B. Ena/VASP Proteins: Regulators of the Actin Cytoskeleton and Cell Migration. Annu. Rev. Cell Dev. Biol. 19, 541–564 (2003).

15. Hansen, S. D. & Mullins, R. D. VASP is a processive actin polymerase that requires monomeric actin for barbed end association. J. Cell Biol. 191, 571–584 (2010).

16. Kühnel, K. et al. The VASP tetramerization domain is a right-handed coiled coil based on a 15-residue repeat. Proc. Natl. Acad. Sci. 101, 17027–17032 (2004).

17. Isambert, H. et al. Flexibility of Actin Filaments Derived from Thermal Fluctuations: EFFECT OF BOUND NUCLEOTIDE, PHALLOIDIN, AND MUSCLE REGULATORY PROTEINS *. J. Biol. Chem. 270, 11437–11444 (1995).

18. Chandrasekaran, A., Graham, K., Stachowiak, J. C. & Rangamani, P. Kinetic trapping organizes actin filaments within liquid-like protein droplets. Nat. Commun. 15, 3139 (2024).

19. Lafuente, E. M. et al. RIAM, an Ena/VASP and Profilin Ligand, Interacts with Rap1-GTP and Mediates Rap1-Induced Adhesion. Dev. Cell 7, 585–595 (2004).

20. Krause, M. et al. Lamellipodin, an Ena/VASP Ligand, Is Implicated in the Regulation of Lamellipodial Dynamics. Dev. Cell 7, 571–583 (2004).

21. Krause, M. et al. Fyn-Binding Protein (Fyb)/Slp-76–Associated Protein (Slap), Ena/Vasodilator-Stimulated Phosphoprotein (Vasp) Proteins and the Arp2/3 Complex Link T Cell Receptor (Tcr) Signaling to the Actin Cytoskeleton. J. Cell Biol. 149, 181–194 (2000).

22. Drees, B. et al. Characterization of the Interaction between Zyxin and Members of the Ena/Vasodilator-stimulated Phosphoprotein Family of Proteins *. J. Biol. Chem. 275, 22503–22511 (2000).

23. Graham, K., et al. Liquid-like condensates mediate competition between actin branching and bundling. Proc. Natl. Acad. Sci. 121, e2309152121 (2024).

24. Cheng, K. W. & Mullins, R. D. Initiation and disassembly of filopodia tip complexes containing VASP and lamellipodin. Mol. Biol. Cell 31, 2021–2034 (2020).

25. Hansen, S. D. & Mullins, R. D. Lamellipodin promotes actin assembly by clustering Ena/VASP proteins and tethering them to actin filaments. eLife 4, e06585 (2015).

26. Montaño-Rendón, F., et al. PtdIns(3,4)P _2_, Lamellipodin, and VASP Coordinate Cytoskeletal Remodeling during Phagocytic Cup Formation in Macrophages. http://biorxiv.org/lookup/doi/10.1101/2022.03.08.483476 (2022) doi:10.1101/2022.03.08.483476.

27. Dimchev, G. et al. Lamellipodin tunes cell migration by stabilizing protrusions and promoting adhesion formation.

28. Chang, Y.-C., Zhang, H., Brennan, M. L. & Wu, J. Crystal structure of Lamellipodin implicates diverse functions in actin polymerization and Ras signaling. Protein Cell 4, 211–219 (2013).

29. Gorai, S. et al. Mechanistic insights into the phosphatidylinositol binding properties of the pleckstrin homology domain of lamellipodin. Mol. Biosyst. 12, 747–757 (2016).

30. Pokrant, T., et al. Ena/VASP clustering at microspike tips involves lamellipodin but not I-BAR proteins, and absolutely requires unconventional myosin-X. Proc. Natl. Acad. Sci. 120, e2217437120 (2023).

31. Mondal, S. et al. Multivalent interactions between molecular components involved in fast endophilin mediated endocytosis drive protein phase separation. Nat. Commun. 13, 5017 (2022).

32. Hansen, S. D. & Mullins, R. D. VASP is a processive actin polymerase that requires monomeric actin for barbed end association. J. Cell Biol. 191, 571–584 (2010).

33. Pollard, T. D. Actin and Actin-Binding Proteins. Cold Spring Harb. Perspect. Biol. 8, a018226 (2016).

34. Winkelman, J. D., Bilancia, C. G., Peifer, M. & Kovar, D. R. Ena/VASP Enabled is a highly processive actin polymerase tailored to self-assemble parallel-bundled F-actin networks with Fascin. Proc. Natl. Acad. Sci. 111, 4121–4126 (2014).

35. Day, K. J. et al. Liquid-like protein interactions catalyse assembly of endocytic vesicles. Nat. Cell Biol. 23, 366–376 (2021).

36. Tebar, F., Sorkina, T., Sorkin, A., Ericsson, M. & Kirchhausen, T. Eps15 Is a Component of Clathrin-coated Pits and Vesicles and Is Located at the Rim of Coated Pits *. J. Biol. Chem. 271, 28727–28730 (1996).

37. Cupers, P., Haar, E. ter, Boll, W. & Kirchhausen, T. Parallel Dimers and Anti-parallel Tetramers Formed by Epidermal Growth Factor Receptor Pathway Substrate Clone 15 (EPS15) *. J. Biol. Chem. 272, 33430–33434 (1997).

38. Cheng, X., Ullo, M. F. & Case, L. B. Reconstitution of Phase-Separated Signaling Clusters and Actin Polymerization on Supported Lipid Bilayers. Front. Cell Dev. Biol. 10, (2022).

39. Li, P. et al. Phase transitions in the assembly of multivalent signalling proteins. Nature 483, 336–340 (2012).

40. Tsukita, K. et al. Phase separation of an actin nucleator by junctional microtubules regulates epithelial function. Sci. Adv. 9, eadf6358 (2023).

41. Yang, S. et al. Self-construction of actin networks through phase separation–induced abLIM1 condensates. Proc. Natl. Acad. Sci. 119, e2122420119 (2022).

42. Yu, Y. & Yoshimura, S. H. Self-assembly of CIP4 drives actin-mediated asymmetric pit-closing in clathrin-mediated endocytosis. Nat. Commun. 14, 4602 (2023).

43. Romero, P. et al. Sequence complexity of disordered protein. Proteins Struct. Funct. Bioinforma. 42, 38–48 (2001).

44. Sun, J. et al. Precise prediction of phase-separation key residues by machine learning. Nat. Commun. 15, 2662 (2024).

45. Mitrea, D. M. et al. Self-interaction of NPM1 modulates multiple mechanisms of liquid–liquid phase separation. Nat. Commun. 9, 842 (2018).

46. Molliex, A. et al. Phase Separation by Low Complexity Domains Promotes Stress Granule Assembly and Drives Pathological Fibrillization. Cell 163, 123–133 (2015).

47. Uversky, V. N. Intrinsically Disordered Proteins and Their Environment: Effects of Strong Denaturants, Temperature, pH, Counter Ions, Membranes, Binding Partners, Osmolytes, and Macromolecular Crowding. Protein J. 28, 305–325 (2009).

48. Brangwynne, C. P., Tompa, P. & Pappu, R. V. Polymer physics of intracellular phase transitions. Nat. Phys. 11, 899–904 (2015).

49. Bachmann, C., Fischer, L., Walter, U. & Reinhard, M. The EVH2 Domain of the Vasodilator-stimulated Phosphoprotein Mediates Tetramerization, F-actin Binding, and Actin Bundle Formation *. J. Biol. Chem. 274, 23549–23557 (1999).

50. Morton, W. M., Ayscough, K. R. & McLaughlin, P. J. Latrunculin alters the actin-monomer subunit interface to prevent polymerization. Nat. Cell Biol. 2, 376–378 (2000).

51. Banani, S. F., Lee, H. O., Hyman, A. A. & Rosen, M. K. Biomolecular condensates: organizers of cellular biochemistry. Nat. Rev. Mol. Cell Biol. 18, 285–298 (2017).

52. Hyman, A. A., Weber, C. A. & Jülicher, F. Liquid-Liquid Phase Separation in Biology. Annu. Rev. Cell Dev. Biol. 30, 39–58 (2014).

53. Burke, K. A., Janke, A. M., Rhine, C. L. & Fawzi, N. L. Residue-by-residue view of in vitro FUS granules that bind the C-terminal domain of RNA polymerase II. Mol. Cell 60, 231–241 (2015).

54. Benmerah, A. et al. AP-2/Eps15 Interaction Is Required for Receptor-mediated Endocytosis. J. Cell Biol. 140, 1055–1062 (1998).

55. Fazioli, F., Minichiello, L., Maťoškovā, B., Wong, W. T. & Di Fiore, P. P. eps15, A Novel Tyrosine Kinase Substrate, Exhibits Transforming Activity. Mol. Cell. Biol. 13, 5814–5828 (1993).

56. Henne, W. M. et al. FCHo Proteins Are Nucleators of Clathrin-Mediated Endocytosis. Science 328, 1281–1284 (2010).

57. Riedl, J. et al. Lifeact: a versatile marker to visualize F-actin. Nat. Methods 5, 605–607 (2008).

58. Schachtner, H. et al. Tissue inducible Lifeact expression allows visualization of actin dynamics in vivo and ex vivo. Eur. J. Cell Biol. 91, 923–929 (2012).

59. Belyy, A., Merino, F., Sitsel, O. & Raunser, S. Structure of the Lifeact–F-actin complex. PLOS Biol. 18, e3000925 (2020).

60. Courtemanche, N., Pollard, T. D. & Chen, Q. Avoiding artifacts when counting polymerized actin in live cells with Lifeact-fluorescent fusion proteins. Nat. Cell Biol. 18, 676–683 (2016).

61. Kang, H. et al. Identification of cation-binding sites on actin that drive polymerization and modulate bending stiffness. Proc. Natl. Acad. Sci. U. S. A. 109, 16923–16927 (2012).

62. Selden, L. A., Gershman, L. C. & Estes, J. E. A kinetic comparison between Mg-actin and Ca-actin. J. Muscle Res. Cell Motil. 7, 215–224 (1986).

63. Limozin, L., Bärmann, M. & Sackmann, E. On the organization of self-assembled actin networks in giant vesicles. Eur. Phys. J. E 10, 319–330 (2003).

64. Miyazaki, M., Chiba, M., Eguchi, H., Ohki, T. & Ishiwata, S. Cell-sized spherical confinement induces the spontaneous formation of contractile actomyosin rings in vitro. Nat. Cell Biol. 17, 480–489 (2015).

65. Claessens, M. M. a. E., Tharmann, R., Kroy, K. & Bausch, A. R. Microstructure and viscoelasticity of confined semiflexible polymer networks. Nat. Phys. 2, 186–189 (2006).

66. Winder, S. J. & Ayscough, K. R. Actin-binding proteins. J. Cell Sci. 118, 651–654 (2005).

67. Holt, M. R. & Koffer, A. Cell motility: proline-rich proteins promote protrusions. Trends Cell Biol. 11, 38– 46 (2001).

68. Hwang, T. et al. Native proline-rich motifs exploit sequence context to target actin-remodeling Ena/VASP protein ENAH. eLife 11, e70680 (2022).

69. Case, L. B., Ditlev, J. A. & Rosen, M. K. Regulation of Transmembrane Signaling by Phase Separation. Annu. Rev. Biophys. 48, 465–494 (2019).

