## Supplementary Information for "Liquid-like condensates that bind actin drive filament polymerization and bundling"

**Affiliations:** 1. Biomedical Engineering, The University of Texas at Austin, Austin, TX, United States. 2. Department of Mechanical and Aerospace Engineering, University of California San Diego, La Jolla, CA, United States. 3. Cell and Molecular Biology, The University of Texas at Austin, Austin, TX, United States. 4. Department of Biochemistry and Structural Biology, University of Texas Health Science Center at San Antonio, San Antonio, TX, United States. 5. Chemical Engineering, The University of Texas at Austin, Austin, TX, United States.

[§]These authors contributed equally

### Supplementary Material

#### Supplementary Tables

**Supplementary Table 1: Table of parameters required to set up the actin model in Cytosim. This set of parameters is common to all the simulations conducted for this study.**

| Parameter | Value | Notes/Reference |
|---|---|---|
| Total time | 600 s | |
| Time step | 0.002 s | |
| Condensate viscosity | 0.5 pN/s μm$^2$ | 50x water |
| **Boundary** | | |
| Shape | Sphere | |
| Radius | 1 μm | |
| Boundary repulsion stiffness | 200 pN/μm for actin filaments;<br>100 pN/μm for crosslinking molecules | This specifies the spring stiffness that acts on the discretized points of each actin filament and crosslinking molecule if the point lies outside the specified boundary. The force on each point is dependent on the distance that it lies beyond the confines of the boundary. |
| **Actin filaments** | | |
| Segmentation length $L_{seg}$ | 0.1 μm (100 nm) | |
| Maximum length | 2πR μm | |
| Polymerization rate $k_{grow}$ | 0.0103 μm/s | Only plus (+) end extension is allowed. This rate is calculated by assuming a final filament length of 2π μm at 600 s. |
| Brownian ratchet force for polymerization | 10 pN | [1] |

| | | |
|---|---|---|
| Actin flexural rigidity $k_{bend}$ | 0.075 pN μm$^2$ | [2] |
| Actin steric repulsion $k_{steric}$ | Radius 3.5 nm<br>Stiffness 1.0 pN/μm | Chosen to ensure the observation of ring structures within the kinetic parameters used in this study as determined from a previous study.[3] |
| **Bivalent crosslinkers (mini-Lpd)** | | |
| Radius | 30 nm | |
| Diffusion rate | 10 μm$^2$/s | |
| Concentration of dimers | 0.40 μM [1000 dimers] | |
| Actin-binding rate<br>(Ring Conditions) | 10.0 (1/s) | Determined from VASP and mini-Lpd simulations. |
| Actin-binding rate<br>(Shell Conditions) | 0.1 (1/s) | Determined from VASP and mini-Lpd simulations. |
| Actin-binding distance | 30 nm | |
| Actin-binding valency | 2 | Each spherical molecule approximates a mini-Lpd dimer. |
| Zero-force actin-unbinding rate (Ring and Shell Conditions) | 1.0 (1/s) | Determined from VASP and mini-Lpd simulations. |
| Actin-unbinding force | 10 pN | Typical values for passive crosslinkers.[4] |
| mini-Lpd steric repulsion | Radius 30 nm<br>Stiffness 1.0 pN/μm | Chosen to ensure the observation of ring structures within the kinetic parameters used in this study as determined from a previous study on tetramers.[3] |
| **Tetravalent crosslinkers (VASP)** | | |
| Radius | 30 nm | |

| | | |
|---|---|---|
| Diffusion rate | 10 μm²/s | |
| Concentration of tetramers | 0.40 μM [1000 tetramers] | |
| Actin-binding rate (Ring Conditions) | 10.0 (1/s) | Determined from VASP and mini-Lpd simulations. |
| Actin-binding rate (Shell Conditions) | 0.1 (1/s) | Determined from VASP and mini-Lpd simulations. |
| Actin-binding distance | 30 nm | |
| Actin-binding valency | 4 | Each spherical molecule approximates a VASP tetramer. |
| Zero-force actin-unbinding rate (Ring and Shell Conditions) | 1.0 (1/s) | Determined from VASP and mini-Lpd simulations. |
| Actin-unbinding force | 10 pN | Typical values for passive crosslinkers.[4] |
| VASP steric repulsion | Radius 30 nm<br>Stiffness 10 pN/μm | Chosen to ensure the observation of ring structures within the kinetic parameters used in this study as determined from a previous study on tetramers.[3] |
| **Monovalent actin binders for dynamic dimerization (mini-Lpd monomers)** | | |
| Radius | 30 nm | |
| Diffusion rate | 10 μm²/s | |
| Concentration of monomers | 0.79 μM [2000 monomers] | Simulations contain 2000 monomers which can form up to a maximum of 1000 bivalent crosslinkers. |
| Actin-binding rate (Ring Conditions) | 10.0 (1/s) | Determined from VASP and mini-Lpd simulations. |
| Actin-binding rate (Shell Conditions) | 0.1 (1/s) | Determined from VASP and mini-Lpd simulations. |

| | | |
|---|---|---|
| Actin-binding distance | 30 nm | |
| Actin-binding valency | 1 | Each spherical molecule approximates a mini-Lpd monomer. |
| Zero-force actin-unbinding rate<br>(Ring and Shell Conditions) | 1.0 (1/s) | Determined from VASP and mini-Lpd simulations. |
| Actin-unbinding force | 10 pN | Typical values for passive crosslinkers.[4] |
| mini-Lpd monomer-actin steric repulsion | Radius $R_{solid}$ = 30 nm<br>Stiffness 10 pN/μm | Chosen to ensure the observation of ring structures within the kinetic parameters used in this study as determined from a previous study on tetramers.[3] |
| Dimerization distance | 90 nm ($3R_{solid}$) | This is the distance between the binding site on solid A and the center of solid B. So, if two solids are in contact, depending on the position of the binding site this distance can scale between $R_{solid}$ and $3R_{solid}$, where $R_{solid}$ is radius of solid. |
| Dimerization splitting force | 10 pN | Used in this study |
| mini-Lpd monomer-monomer steric repulsion | Radius 30 nm<br>Stiffness 5.0 pN/μm | Chosen empirically to ensure adequate dimerization reactions occur to support the observation of ring structures in Figure 5. |

**Supplementary Table 2: Additional parameters employed to simulate specific simulations discussed in this paper**

| Parameter | Value | Notes/Reference |
|---|---|---|
| **Parameters for simulations with mixtures of bivalent and tetravalent crosslinkers (Figures S2 and S3)** | | |
| Tetravalent: Bivalent crosslinker copy number ratios | {1000:0, 750:250, 500:500, 250:750, 0:1000} | Used in this study |
| Tetravalent-Bivalent crosslinker copy number kinetic conditions | {Ring-Ring, Ring-Shell, Shell-Ring} | Kinetic conditions considered, values shown in Table S1. |
| **Parameters for simulations with bivalent crosslinkers (Figure 4)** | | |
| Binding rates $k_{bind}^{mini-Lpd}$ | {$10^{-3}$, $10^{-2}$, $10^{-1}$, $10^{0}$, $10^{+1}$} (1/s) | Range determined based on a previous study with tetramers.[3] |
| Zero-force actin unbinding rates $k_{unbind}^{mini-Lpd}$ | {$10^{-3}$, $10^{-2}$, $10^{-1}$, $10^{0}$, $10^{+1}$} (1/s) | |
| **Parameters for simulations of dynamically dimerizing proteins (Figure 5)** | | |
| Dimer forming rates $k_{dimer\ form}$ | {$10^{-3}$, $10^{-2}$, $10^{-1}$, $10^{0}$, $10^{+1}$} (1/s) | Used in this study |
| Dimer splitting rates $k_{dimer\ split}$ | {$10^{-3}$, $10^{-2}$, $10^{-1}$, $10^{0}$, $10^{+1}$} (1/s) | Used in this study |

#### Supplementary Methods

##### Chemical and mechanical framework employed in Cytosim

Simulations were performed in Cytosim (https://gitlab.com/f-nedelec/cytosim), an agent-based modeling framework which simulates the chemical dynamics and mechanical properties of filament networks. Cytosim models filament dynamics and diffusing species by numerically solving a constrained Langevin framework in a viscous medium at short time intervals. Actin filaments are represented as inextensible fibers composed of a series of linear segments of length 100 nm connected at hinge points to allow for bending. Cytosim computes the bending energy of the fiber using the specified flexural rigidity in the input parameters. In this study, cross linking molecules (mini-Lpd, VASP, mini-Lpd monomers) are modeled as spherical solids of radius 30 nm with a specified number and type of binding sites corresponding with the class of molecule (**Fig. S1**). The binding distance specifies the radius within which the concentration of the corresponding reactant is considered as part of the binding reaction. The unbinding rate specifies the rate constant used in a Bell's law model representation of slip bond unbinding kinetics. We started with the simulation framework in [3,5]. We assumed that only a subset of crosslinking molecules participate in bundling owing to steric accessibility issues. The condensate was represented by a spherical volume of radius 1.0 μm with a rigid, repulsive boundary. We considered 30 actin filaments within the condensate, each of length 0.1 μm. The actin filament elongation rate is 0.0103 μm/s, calculated assuming a final filament length of 2π μm. The simulation run time is 600 s and was informed by experiments. Please refer to Supplementary Table S1 for a detailed description of the parameters used in the model and Supplementary Table S2 for a detailed description of simulation specific parameters varied. In this study, we performed simulations by modeling mini-Lpd molecules as solids with two binding sites (**Fig. S2** and **Fig. 4**) and VASP molecules as solids with four actin-binding sites (**Fig. S2**). Additionally, we also modified the codebase to model mini-Lpd molecules as those that dimerize based on a given formation and splitting rates.

##### Position evolution

Cytosim uses the Langevin equation to calculate the evolution of discretized points over time, thus describing the 3D position of each actin filament and crosslinking molecule in the system at each time step for the duration of the simulation. In a 3D system of N particles, there are a total of 3N coordinates where each particle i has its coordinates given by $x_i = \{x_{i1}, x_{i2}, x_{i3}\}$. Each particle's position $x_i$ is then evolved along each dimension j as governed by the following stochastic differential equation:

$$dx^{ij}(t) = \mu f_{tot}^{ij}(t)dt + dB_j(t)$$

Here, μ is the viscosity of the solvent, $f^{ij}_{tot}(t)$ is the total force acting on each particle as a function of time, and $B_j(t)$ is the diffusion (noise) term. The noise term is given by a randomly sampled variable from a normal distribution centered around a mean of 0 with a standard deviation of $\sqrt{2D^i dt}$. The diffusion constant D is given by the Einstein relation $D = \mu k_B T$ where $k_B$ is the Boltzmann constant and T is temperature.

**Steric considerations**

It is important to consider steric repulsion potentials to prevent spatial overlap of molecules. Additionally, crowding also affects the effective mobility of molecules thereby altering the propensities of chemical reactions in our system. As such, we employ a steric repulsion potential between the diffusing elements in our simulations, the crosslinking molecules and actin filaments.

**Dynamic dimerization model**

The dynamic dimerization model simulates independently diffusing monomers as solids that each have a single actin binding site and a single dimerization site capable of binding to another monomer to dynamically form dimers. This introduces new input parameters that describe the kinetics of forming and splitting dimers. This differs from the other simulations where bivalent crosslinkers are prescribed as a single solid representing a static dimer with two binding sites. The implementation of our dynamic dimerization model required edits to the Cytosim source code to simulate dimerization reactions. These source code edits are available on the GitHub repository available with this publication.

**Actin-covered surface area calculation**

At the end of the simulation time, each actin filament grows to approach the length of the circumference of the spherical condensate. At this point, each actin filament will lie primarily at or near the condensate surface (defined with a threshold distance from the boundary of 100 nm). By discretizing the surface of the condensate to an icosphere and each actin filament into discrete monomers, the surface density at each timepoint in our simulation is obtained via the fraction of occupied triangles on the icosphere. The icosphere was generated by dividing the initial eight triangles 3 more times to generate triangles whose effective size was comparable to 10 actin monomers.[3]

#### Supporting Figures

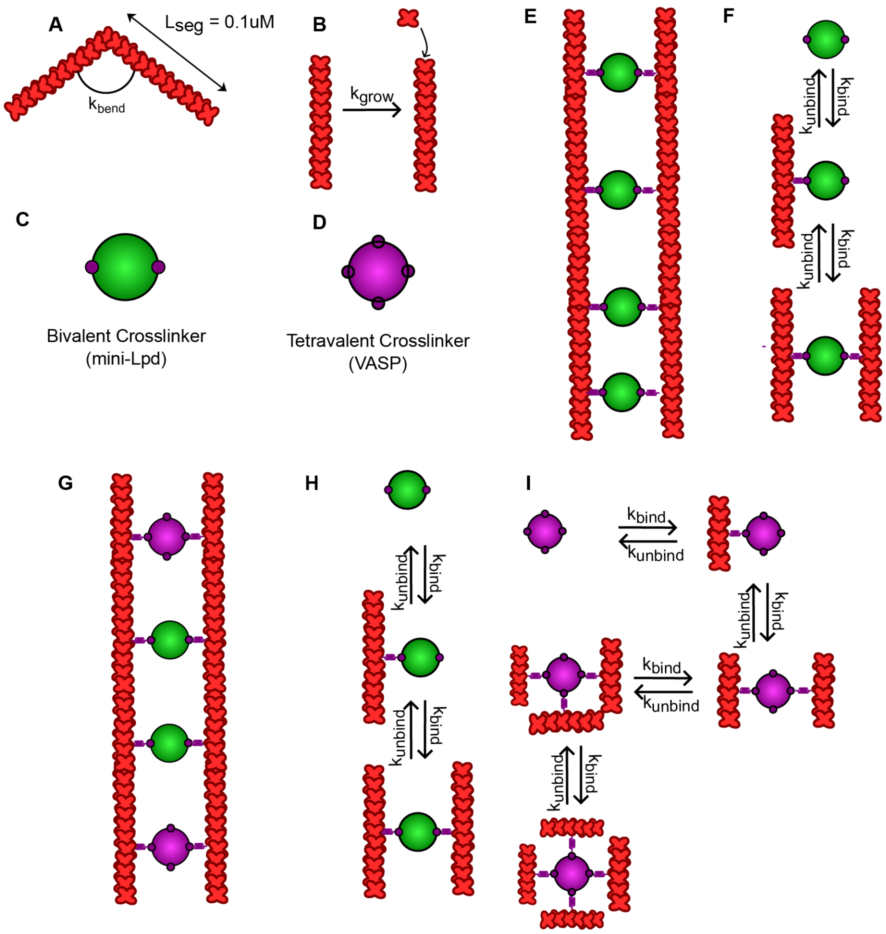


**Figure S1: Schematic representation of chemical species and reactions considered. A.** Actin filaments are modeled as inextensible segments of length $L_{seg}$ that can bend along hinge points based on flexural rigidity $k_{bend}$. **B.** Filaments in these simulations are allowed to grow deterministically at rate $k_{grow}$ = 0.0103 μm/s. **C.** Bivalent crosslinkers mimic mini-Lpd dimers and are represented as green-colored solids that have two binding sites (small, purple circles) which can bind and crosslink actin filament. **D.** Tetravalent crosslinkers mimic VASP (in purple) and have four actin-binding sites. **E.** Cartoon representation of two filaments crosslinked by mini-Lpd. **F.** Each of the two actin binding domains in a dimeric crosslinker bind actin at rate $k_{bind}$ and unbind in a force-sensitive manner with unbinding rate $k_{unbind}$. **G.** Cartoon representation of two actin filaments crosslinked by both mini-Lpd and VASP molecules. **H, I.** For simulations with mixtures of mini-Lpd and VASP, the crosslinking reactions considered are shown. Please refer to Supporting Information, Tables S1 and S2 for more information on model parameters used in this study.

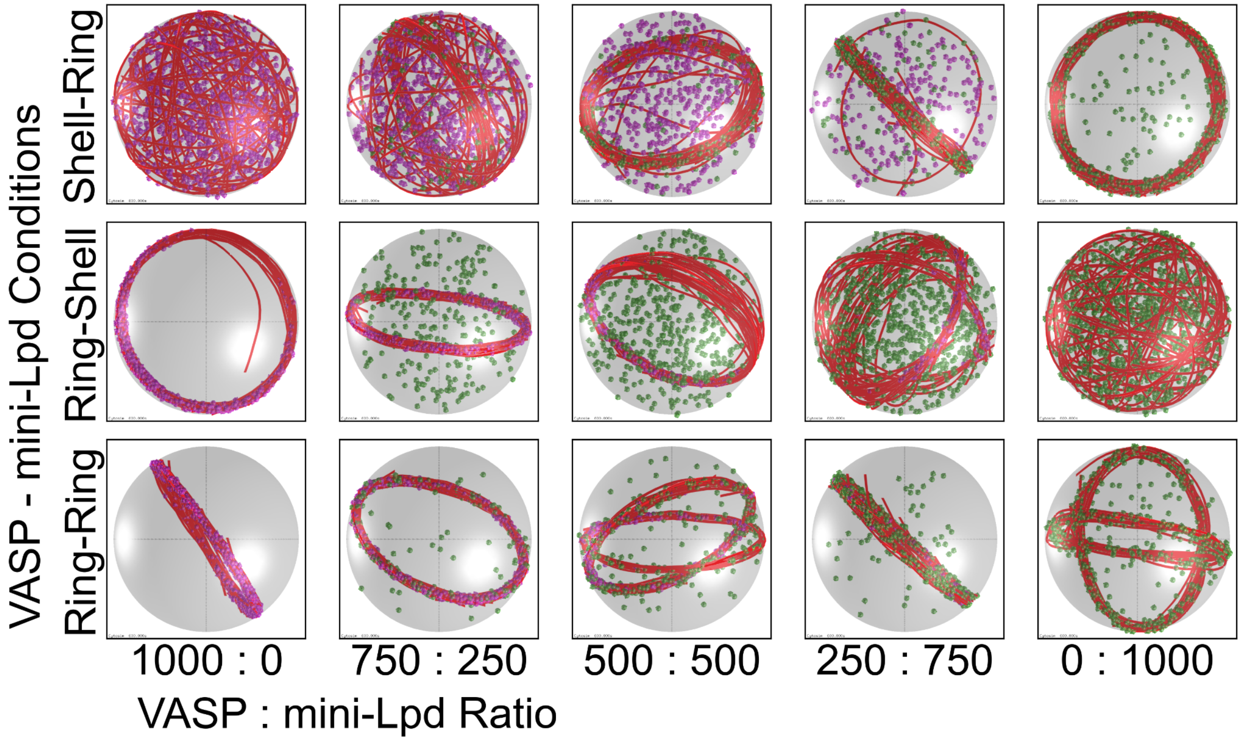


**Figure S2: Simulations show that ring and shell-shaped actin network structures are conserved within condensates of mini-Lpd and VASP of various mole ratios.** Representative final snapshots (t = 600 s) from simulations for three different crosslinker kinetic conditions at various VASP to mini-Lpd mole ratios. Condensates are spherical (R = 1 μm), contain 30 actin filaments (red), and a total of 1000 crosslinkers (tetravalent crosslinkers in purple and bivalent crosslinkers in green). Please refer to the Supplementary Methods section for a detailed description of the model. Ring-forming kinetics ($k_{bind}$ = 10.0 $s^{-1}$, $k_{unbind}$ = 1.0 $s^{-1}$) or shell-forming kinetics ($k_{bind}$ = 0.1 $s^{-1}$, $k_{unbind}$ = 1.0 $s^{-1}$) are the same for both types of crosslinkers. The polymerization rate at the plus (+) end is constant at 0.0103 μm/s, and neither end undergoes depolymerization. See **Supplementary Movie M1** for a video of representative trajectories.

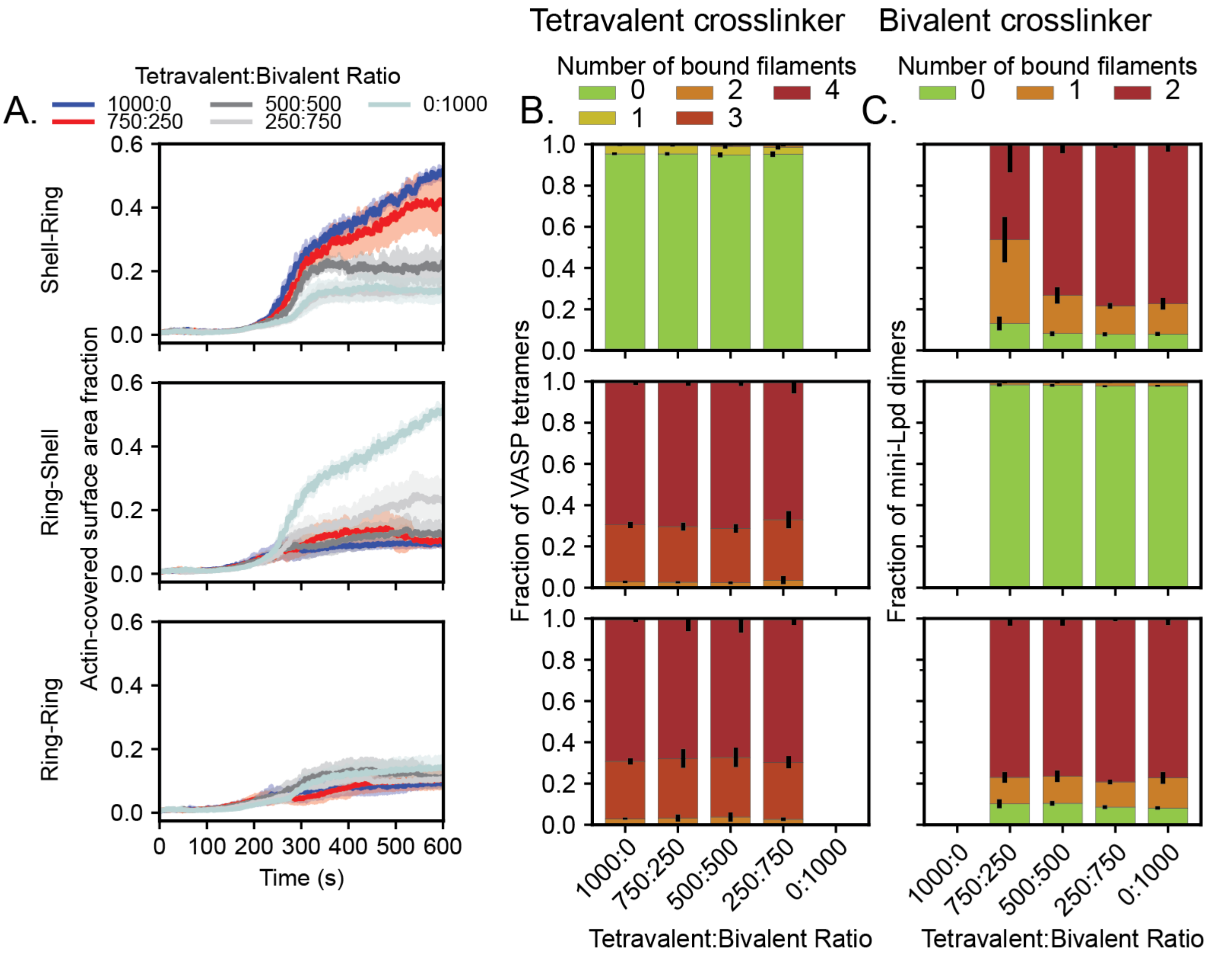


**Figure S3. Analysis metrics used to understand actin organization within condensates at various number ratios of tetravalent and bivalent crosslinkers. A.** Mean (solid line) and standard deviation (shaded area) of the condensate surface area covered with actin is shown as a time series. Time series is colored by tetravalent: bivalent ratio. Each subpanel shows simulations at different actin binding parameters used. For example, Shell-Ring corresponds to kinetic parameter choice where we have observed shell formation with 1000 molecules of tetravalent crosslinker and ring formation when simulated with 1000 molecules of bivalent crosslinker respectively. **B.** The stacked bar graphs show the distribution of tetravalent crosslinkers (VASP) molecules among various allowed valency states (mentioned above) as we change the copy number ratio. Error bars represent standard deviation. **C.** The stacked bar graphs show the distribution of bivalent crosslinkers (mini-Lpd) molecules among various allowed valency states (mentioned above) as we change the copy number ratio. Error bars represent standard deviation. Data used: 5 replicates per kinetic condition (namely, Shell-Ring, Ring-shell, and Ring-Ring), bar graphs generated with data from the last 30 snapshots from each of the replicates.

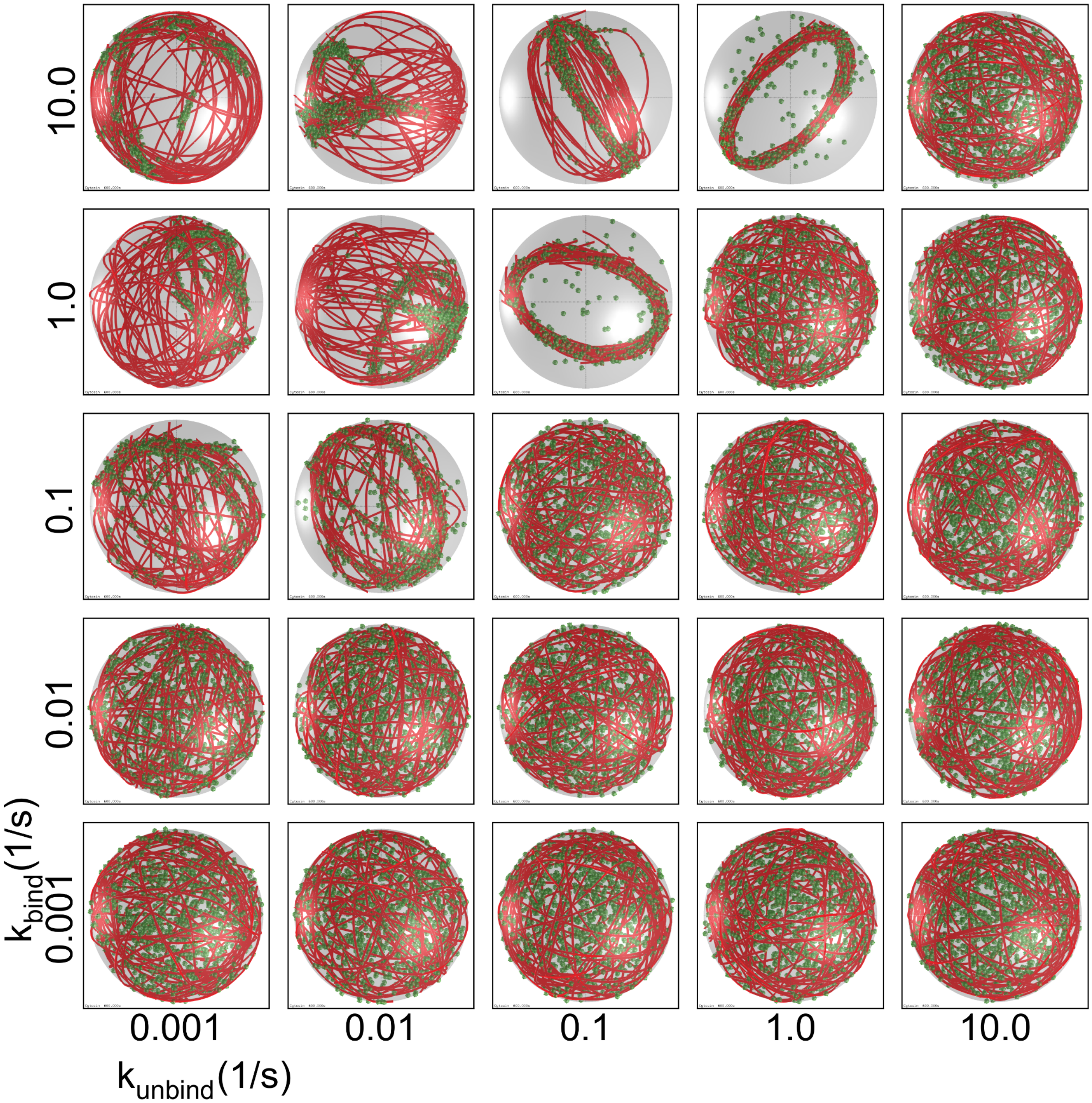


**Figure S4: Simulations show that mini-Lpd kinetics affect actin network organization in liquid-liquid phase-separated mini-Lpd dimer condensates.** Representative final snapshots (t = 600 s) from simulations at various binding and unbinding rates within spherical condensates (R = 1 μm) containing 30 actin filaments (red) and 1000 bivalent crosslinkers (green). Please refer to the Supplementary Methods section for a detailed description of the model. The binding rates of the bivalent crosslinkers are varied along each column, and unbinding rates are varied along each row. The polymerization rate at the plus (+) end is constant at 0.0103 μm/s, and neither end undergoes depolymerization. Also see **Supplementary Movie M3**.

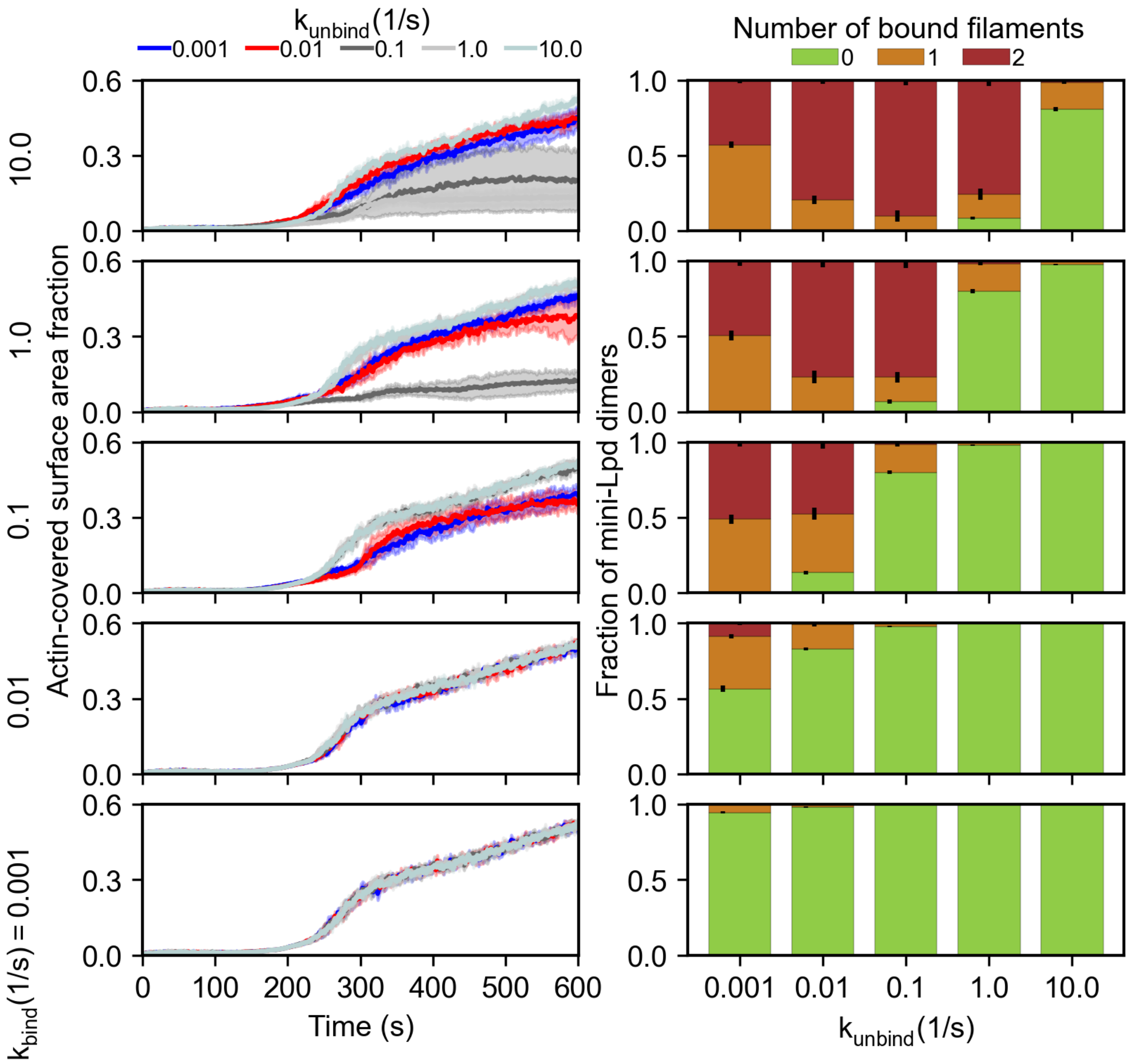


**Figure S5: Kinetics of the actin-covered surface area fraction time series and the fraction of mini-Lpd bound to zero, one, and two actin filaments.** The mini-Lpd binding rate is changed across the panels (shown on the left side of the figure). **A.** The mini-Lpd unbinding rate is varied and displayed as a time series of the actin-covered surface area fraction. **B.** Stacked bar graphs representing the fraction of bivalent crosslinkers bound to 0, 1, or 2 actin filaments for each condition. The error bars represent the standard deviation. Ten replicates are considered per condition, and the data was obtained from the last 30 snapshots (5%) of each replicate.

**References:**


1. Mogilner, A., and Oster, G. (1996). Cell motility driven by actin polymerization. Biophys. J. *71*, 3030–3045.

2. Gittes, F., Mickey, B., Nettleton, J., and Howard, J. (1993). Flexural rigidity of microtubules and actin filaments measured from thermal fluctuations in shape. J. Cell Biol. *120*, 923–934.

3. Chandrasekaran, A., Graham, K., Stachowiak, J.C., and Rangamani, P. (2024). Kinetic trapping organizes actin filaments within liquid-like protein droplets. Nat. Commun. *15*, 3139.

4. Ferrer, J.M., Lee, H., Chen, J., Pelz, B., Nakamura, F., Kamm, R.D., and Lang, M.J. (2008). Measuring molecular rupture forces between single actin filaments and actin-binding proteins. Proc. Natl. Acad. Sci. U. S. A. *105*, 9221–9226.

5. Graham, K., Chandrasekaran, A., Wang, L., Yang, N., Lafer, E.M., Rangamani, P., and Stachowiak, J.C. (2024). Liquid-like condensates mediate competition between actin branching and bundling. Proc. Natl. Acad. Sci. U. S. A. *121*, e2309152121.

**Title:** Supplementary Movie 1.
**Description:** Movie gallery depicting representative trajectories from simulations of three different crosslinker kinetic conditions at various VASP to mini-Lpd mole ratios. Condensates are spherical (R = 1 μm), contain 30 actin filaments (red), and a total of 1000 crosslinkers (tetravalent crosslinkers in purple and bivalent crosslinkers in green). Ring-forming kinetics ($k_{bind}$ = 10.0 $s^{-1}$, $k_{unbind}$ = 1.0 $s^{-1}$) or shell-forming kinetics ($k_{bind}$ = 0.1 $s^{-1}$, $k_{unbind}$ = 1.0 $s^{-1}$) are the same for both types of crosslinkers. The polymerization rate at the plus (+) end is constant at 0.0103 μm/s, and neither end undergoes depolymerization. $T_{sim}$ = 600 s, $\Delta t_{frame}$ = 5 s

**Title:** Supplementary Movie 2.
**Description:** Movie gallery depicting representative trajectories from simulations of spherical condensates (R = 1 μm) containing 30 actin filaments (red) and 1000 bivalent crosslinkers (green spheres), representative of mini-Lpd dimers, at $k_{bind}$ (y-axis) and $k_{unbind}$ (x-axis) values varied over 3 orders of magnitude. The binding rates of the bivalent crosslinkers are varied along each column, and unbinding rates are varied along each row. The polymerization rate at the plus (+) end is constant at 0.0103 μm/s, and neither end undergoes depolymerization. Structures observed in wider parameter sweeps are captured by this reduced size gallery. $T_{sim}$ = 600 s, $\Delta t_{frame}$ = 5 s

**Title:** Supplementary Movie 3.
**Description:** Movie gallery depicting representative trajectories from simulations of spherical condensates (R = 1 μm) containing of 30 actin filaments (red) and 1000 bivalent crosslinkers (green spheres), representative of mini-Lpd dimers, at $k_{bind}$ (y-axis) and $k_{unbind}$ (x-axis) values varied over 5 orders of magnitude. The binding rates of the bivalent crosslinkers are varied along each column, and unbinding rates are varied along each row. The polymerization rate at the plus (+) end is constant at 0.0103 μm/s, and neither end undergoes depolymerization. $T_{sim}$ = 600 s, $\Delta t_{frame}$ = 5 s

**Title:** Supplementary Movie 4.
**Description:** Movie gallery depicting representative trajectories from simulations of spherical condensates (R = 1 μm) containing 30 actin filaments (red) and 2000 dynamically dimerizing monomers (green spheres), representative of mini-Lpd monomers, at various $k_{dimer\ form}$ (y-axis) and $k_{dimer\ split}$ (x-axis) values. The dimer formation rates of the monomers are varied along each column, and dimer splitting rates are varied along each row. Actin-binding kinetics were chosen from previous simulations (**Fig. 4**) to correspond to ring-forming conditions for bivalent crosslinkers. The polymerization rate at the plus (+) end is constant at 0.0103 μm/s, and neither end undergoes depolymerization. $T_{sim}$ = 600 s, $\Delta t_{frame}$ = 5 s
